# Hippo effector, Yorkie, is a Tumor Suppressor in Select *Drosophila* Squamous Epithelia

**DOI:** 10.1101/2023.10.15.562319

**Authors:** Rachita Bhattacharya, Jaya Kumari, Shweta Banerjee, Jyoti Tripathi, Nitin Mohan, Pradip Sinha

## Abstract

Out-of-context gain of nuclear signaling of mammalian YAP/TAZ or *Drosophila* Yki—the transcription cofactors of the highly conserved Hippo tumor suppressor pathway—is oncogenic. By contrast, in mechanically strained squamous epithelia (SE), YAP/TAZ/Yki displays developmentally programmed nuclear translocation, leading to its constitutive signaling. How organ homeostasis is maintained in constitutively YAP/TAZ/Yki signaling SE is unclear. Here, we show that Yki signaling negatively regulates the cell growth-promoting PI3K/Akt/TOR signaling in the SEs in the tubular organs of *Drosophila*. Thus, in the adult male accessory gland (MAG), knockdown of Yki signaling upregulates PI3K/Akt/TOR signaling in its SE-lined lumen, inducing cell hypertrophy, culminating in squamous cell carcinoma (SCC). MAG SCC-bearing adults display early mortality due to cancer cachexia, which is reversed by simultaneous knockdown of a secreted factor, ImpL2—a *Drosophila* homolog of mammalian IGFBP7—without arresting tumor progression *per se*. By contrast, a knockdown of PI3K/Akt/TOR signaling suppresses MAG SCC, reversing adult mortality. In the SE-lined lumens in other tubular organs, like the dorsal trunk of larval tracheal airways or adult Malpighian tubules, too, knockdown of Yki signaling triggers PI3K/Akt/TOR-induced cell hypertrophy and loss of epithelial homeostasis, culminating in their tumor-like transformation. Thus, Yki signaling turns tumor suppressive in the SEs of tubular organs in *Drosophila* by arresting runaway PI3K/Akt/TOR signaling.

## INTRODUCTION

Each organ in a metazoan grows predictably and stops cell proliferation after reaching a predefined size/cell number under the regulation of organ-intrinsic growth control mechanisms, such as the highly conserved Hippo signaling pathway (Sebe-Pedrs et al., 2012) (for review, see (Ma et al., 2018; Pan, 2022; Yu et al., 2015)). The discovery and unfolding of this pathway traces back to the identification of an atypical cadherin, Fat (Bryant et al., 1988; Cho et al., 2006; Mahoney et al., 1991), in *Drosophila*, as the most upstream component of intrinsic growth control pathway that restrains cell proliferation to predetermined organ sizes (Bryant and Levinson, 1985; Bryant and Simpson, 1984; Bryant et al., 1988). Tissue overgrowth induced by loss of Fat, individual cytoplasmic kinases of the Hippo pathway, like Warts (Justice et al., 1995; Xu et al., 1995) or Hippo (Harvey et al., 2003) and, alternatively, upon activation of its TEAD-binding transcription cofactor—Yki (Huang et al., 2005) in *Drosophila* and YAP/TAZ in vertebrates (Dong et al., 2007)—were thus construed as proofs of its instructive signaling for organ size regulation (Huang et al., 2005; Willecke et al., 2006) (for review, see (Pan, 2022)). This long-standing interpretation of an instructive role of YAP/TAZ/Yki signaling in organ growth regulation (Ma et al., 2018; Moya and Halder, 2018; Yu et al., 2015; Zheng and Pan, 2019; for review, see Zanconato et al. 2016) has recently been questioned based on two contrarian lines of evidence. First, organ growth is unaffected by a comprehensive genetic abrogation of Yki signaling: for instance, a simultaneous loss of *yki* and its nuclear partner, the TEAD domain transcription factor, Scalloped (Sd). Second, endogenous YAP/TAZ/Yki signaling outputs are not concordant with cell proliferation and growth of a developing organ: for instance, the transcriptional network of YAP/TAZ/Yki remains active long after cessation of endogenous growth in the eye primordium in *Drosophila* or mouse liver (Kowalczyk et al., 2022). Thus, while out-of-context activation of YAP/TAZ/Yki signaling is oncogenic (for review, see (Piccolo et al., 2022; Zheng and Pan, 2019)) or drives cell proliferation during organ regeneration (Oh et al., 2018) (for review, see (Moya and Halder, 2018; Pocaterra et al., 2020; Wang et al., 2018) and wound healing (Tsai et al., 2017) (for review, see (Dey et al., 2020; Lee et al., 2014)), its instructive role in organ size homeostasis, if any, remains elusive (Kowalczyk et al., 2022).

Could Yki/YAP/TAZ signaling be instructive for organ size regulations where they signal perpetually? For instance, squamous epithelia (SE) of all organs examined so far show overt mechanostrain-linked nuclear localization and chronic signaling of YAP/TAZ/Yki (for review, see (Dupont et al., 2011; Halder et al., 2012; Pocaterra et al., 2020). These include mammalian cholangiocytes (Kowalczyk et al., 2022), endothelial cells (Dupont et al., 2011; Hooglugt et al., 2021), stretched fibroblasts (Calvo et al., 2013; Liu et al., 2015), lung alveolar AT1 cells (Li et al., 2018; Mahoney et al., 2014), and stretched retinal pigment epithelial cells of zebrafish (Moreno-Marmol et al., 2018). In *Drosophila,* Yki shows perpetual nuclear signaling in the SE of peripodium of imaginal discs or follicles egg chambers (Borreguero-Muñoz et al., 2019; Fletcher et al., 2018; Friesen and Hariharan, 2023), wherein its loss compromises cell shape and size.

In mechanostrained SE of a tubular organ like mammalian airways (lungs), loss of YAP or TAZ was seen to correlate with squamous cell carcinoma (SCC) (Huang et al., 2017) or development of alveolar cysts (Mahoney et al., 2014; van Soldt et al., 2019). Thus, unlike its oncogenic activation, loss of YAP/TAZ may display tumor suppressor-like function in mechanostrained SEs of tubular organs via unknown mechanisms. To explore such mechanisms, here, we have examined the role of nuclear Yki signaling in the mechanostrained epithelial linings of the lumens of three tubular organs in *Drosophila,* namely, adult male accessory glands (MAG), larval tracheal dorsal trunks, and adult renal Malpighian tubules. We show that the knockdown of Yki signaling upregulates the cell growth-promoting PI3K/Akt/TOR signaling in the MAG SE, triggering cell hypertrophy and lethal, cachexia-inducing SCC. Likewise, in dorsal trunks of larval tracheal airways and adult Malphigian tubules, PI3K/Akt/TOR signaling triggered by *yki* knockdown culminates in hypertrophy and tumor-like transformations of their respective SEs. Our findings reveal that a developmentally programmed chronic Yki signaling—reminiscent of a tumor suppressor— represses runaway PI3K/Akt/TOR signaling and is instructive for organ homeostasis in *Drosophila* tubular organs.

## RESULTS

### Adult MAG displays cell flattening-linked nuclear Yki signaling

Previously, in *Drosophila,* chronic nuclear Yki signaling was shown in the squamous epithelia (SE) of the peripodial membrane of imaginal discs (Fletcher et al., 2018; Friesen and Hariharan, 2023) and ovarian follicles (Borreguero-Muñoz et al., 2019; Fletcher et al., 2018), as a fallout of their mechanical strain. Here, we have further asked if SE of the adult male accessory gland, MAG (Heifetz et al., 2000), too, displays nuclear Yki signaling.

Each of the paired *Drosophila* adult MAG (**Fig. S1**) represents an oblong tube (**Fig. 1A**) that opens into a common ejaculatory duct. Anatomically, MAG SE is distinct from the peripodial or ovarian follicular SEs in that it lines the lumen of a tubular organ. The basolateral membrane of MAG SE marked by Lgl (Grzeschik et al., 2010) rests on a Viking-marked basement membrane, encircled by concentric muscle fibers capable of pulsatile contractions (**Fig. 1A**, also see (Kumari and Sinha, 2021; Rambur et al., 2020)). The pupal MAG (**Fig. 1B**) is a miniaturized version of its adult counterpart (**Fig. 1C**); the apical area of its columnar cells (**Fig. 1B’**) is smaller than its squamous adult counterpart (**Fig. 1C’-D**). A progressive, columnar-to-squamous cell flattening of this epithelial lining of the MAG lumen is achieved by the third day after adult eclosion with accompanying nuclear translocation of the Hippo transcription cofactor, Yki (**Fig. S1**). During this transition, MAG SE displays dilution of an apical membrane determinant, Crumbs (Crb, see (Bulgakova and Knust, 2009; Tepass et al., 1990)) (**Fig. 1E-G**, (Fletcher et al., 2018)) and shortening of the FasIII-marked septate junctions (Baumann, 2001) (**Fig. 1H**). Finally, the nuclear translocation of Yki in the flattened adult MAG SE (Yki::GFP, **Fig. 1I-K**) goes hand in hand with that of its reporter *Diap1*-*lacZ,* a P(*lacZ)* insertion in 5’ UTR of *Diap1* transcription unit that also carries a nuclear localization tag (**Fig. 1L**; also see (Harvey et al., 2003; Zhang et al., 2008). Therefore, in the subsequent part of this study, we have used nuclear translocation of Yki::GFP or Diap1-lacZ as a readout for Yki signaling. These cell flattening-linked hallmarks of MAG are reminiscent of those seen in the adult ovarian follicular and peripodial epithelia (Fletcher et al., 2018; Friesen and Hariharan, 2023; Kowalczyk et al., 2022).

**Fig. 1.**
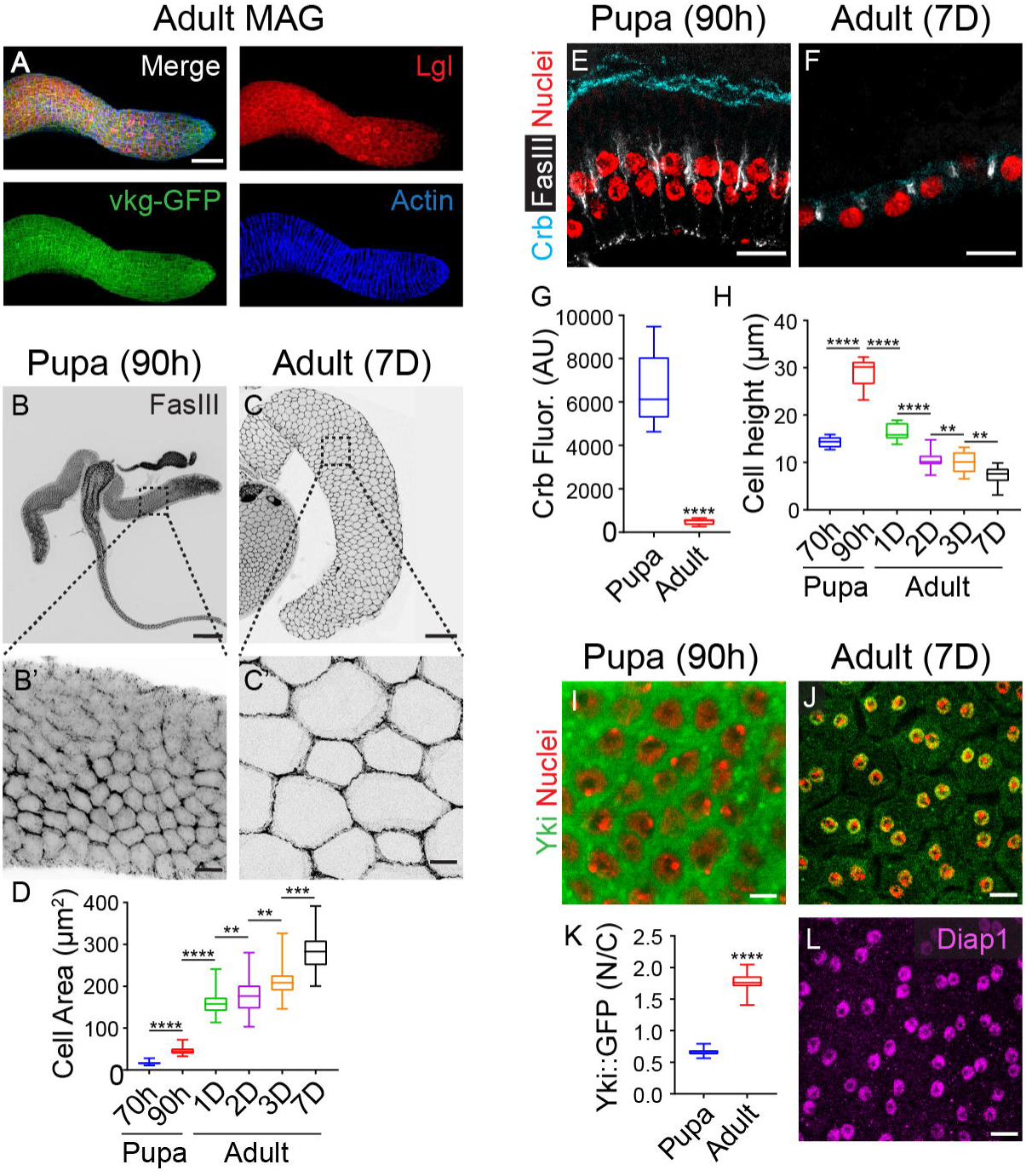
Cell flattening-linked nuclear translocation of Yki in squamous epithelium of adult MAG. (**A**) Maximum intensity projections of optical sections of a MAG from a 7-day-old adult male displaying basolateral membrane (Lgl, red) and basement membrane (*vkg*-*GFP*, green) of its squamous epithelium, wrapped by outer rings of contractile muscle (actin, blue). **(B-C)** Comparison of pupal (B) and adult (C) MAG immunostained for the septate junction marker, FasIII (grey). Magnified boxed areas of these images are displayed in the respective bottom panel (B’, C’). (**D**) Quantifying MAG cell areas (mean ± *SD*, *n*=50 for each stage) from pupae and adults. Unpaired *t-*test for cell area between 70h and 90h pupae, \*\*\*\**p*<0.0001; between 90h pupae and 1D adults, \*\*\*\**p*<0.0001; between 1D and 2D adults, \*\**p*=0.0041; between 2D and 3D adults, \*\*\*\**p*<0.0001, and, finally, between 3D and 7D adults, \*\*\*\**p*<0.0001. **(E-H)** X-Y longitudinal sections of FasIII-stained pupal (E) and adult (F) MAG marked for apical membrane (Crb::GFP, pseudocolored in cyan) and nuclei (TO-PRO, red). Note the striking dilution of the apical membrane in the squamous epithelium of adult MAG by contrast to their columnar pupal counterpart. Quantification of Crb::GFP fluorescence intensity (G). Unpaired *t*-test of average fluorescence intensity between 90h pupae and 7D adults, \*\*\*\**p*<0.0001. Data are represented as mean ± *SD* (*n*=5). Quantification of cell heights (H) at pupal and adult stages of development: unpaired *t*-test of average cell height between 70h and 90h pupae, \*\*\*\**p*<0.0001; between 90h pupae and 1D adults, \*\*\*\**p*<0.0001; between 1D and 2D adults, \*\*\*\**p*=0.0001; between 2D and 3D adults, \*\**p*<0.0041, between 3D and 7D adults, \*\**p*<0.0041; each data point represents mean ± *SD* (*n*=10). **(I-L)** Yki (Yki::GFP, green) is cytoplasmic in pupal (I) and nuclear in adult MAG (J); nuclei are marked by TO-PRO (red). Quantification of nuclear-to-cytoplasmic (N/C) fluorescence of Yki::GFP (*n*=20) (K); \*\*\*\**p*<0.0001. Adult MAG squamous epithelium displaying expression of a Yki target, *Diap1*-*lacZ* (β-gal, magenta) (L). Age of pupal MAG is shown in hours (h) after puparium formation, and those of the adults are shown in days (D) after eclosion. Scale bar: (A-C) 100 µm, (B’, C’, E-L) 10 µm.

Thus, the adult MAG, the squamous epithelial lining, displays nuclear Yki signaling for its entire lifetime.

### Loss of Yki signaling in adult MAG SE induces cell hypertrophy, SCC, and cancer cachexia

To test the fallout of the knockdown of nuclear Yki signaling, we used the *ovulin-Gal4* (*ov-Gal4*) driver (Heifetz et al., 2000). Ovulin is a protein synthesized in MAG and transported to the females during mating; it induces an increase in ovulation by relaxing the oviduct (Heifetz et al., 2000; Herndon and Wolfner, 1995). We noticed that the *ov-Gal4* driver is turned on selectively only in the MAG tubules in 3-day-old adult MAG **(Fig. S1**; also see (Monsma et al., 1990)). However, the expression of *ov-Gal4* progressively diminishes unless these adult males mate with females. Thus, by adding females on the third day after adult eclosion, a robust expression of *ov-Gal4,* coinciding with nuclear Yki signaling (**Fig. S1**), can be sustained for a significant part of the adult male life span (**Fig. S1**).

Previously, it was shown that the knockdown of nuclear Yki signaling in mechanically strained epithelia—like the peripodial layer of imaginal discs or follicular epithelium enveloping the growing oocytes—reduces cell size (Fletcher et al., 2018; Friesen and Hariharan, 2023). By contrast, in MAG SE, loss of Yki signaling induced hypertrophy within two days of Yki signaling knockdown: that is, in a 5-day-old adult (*ov>yki-IR,* **Fig. 2A**). After four days *of yki* knockdown: that is, after seven days (7D) of adult eclosion, the MAG SE showed loss of FasIII-marked septate junctions (**Fig. 2B**), disrupted cytoskeletal architecture (α-tubulin, **Fig. 2C-D**), loss of basement membrane integrity, as revealed by altered distribution of β-integrin (**Fig. 2E-F**, see (Farahani et al., 2014), and expression of matrix metalloprotease (MMPs, **Fig. 2G-H**, (Khan et al., 2013)). These cytoarchitectural perturbations and MMP expression are hallmarks of cancerous epithelial transformation (Aseervatham, 2020; Khan et al., 2013; Rambur et al., 2020). Finally, MAG SCC induced by *yki* downregulation increases nuclear volume, suggesting cancer progression-linked endoreplication (**Fig. 2I-K**, for review, see (Fox and Duronio, 2013; Zhang et al., 2022)). Indeed, endoreplication-linked SCC has been reported earlier in humans (Fox and Duronio, 2013), which, in the context of *Drosophila* MAG, may be primed by its binucleate cell state (Box et al., 2022; Taniguchi et al., 2012).

**Fig. 2.**
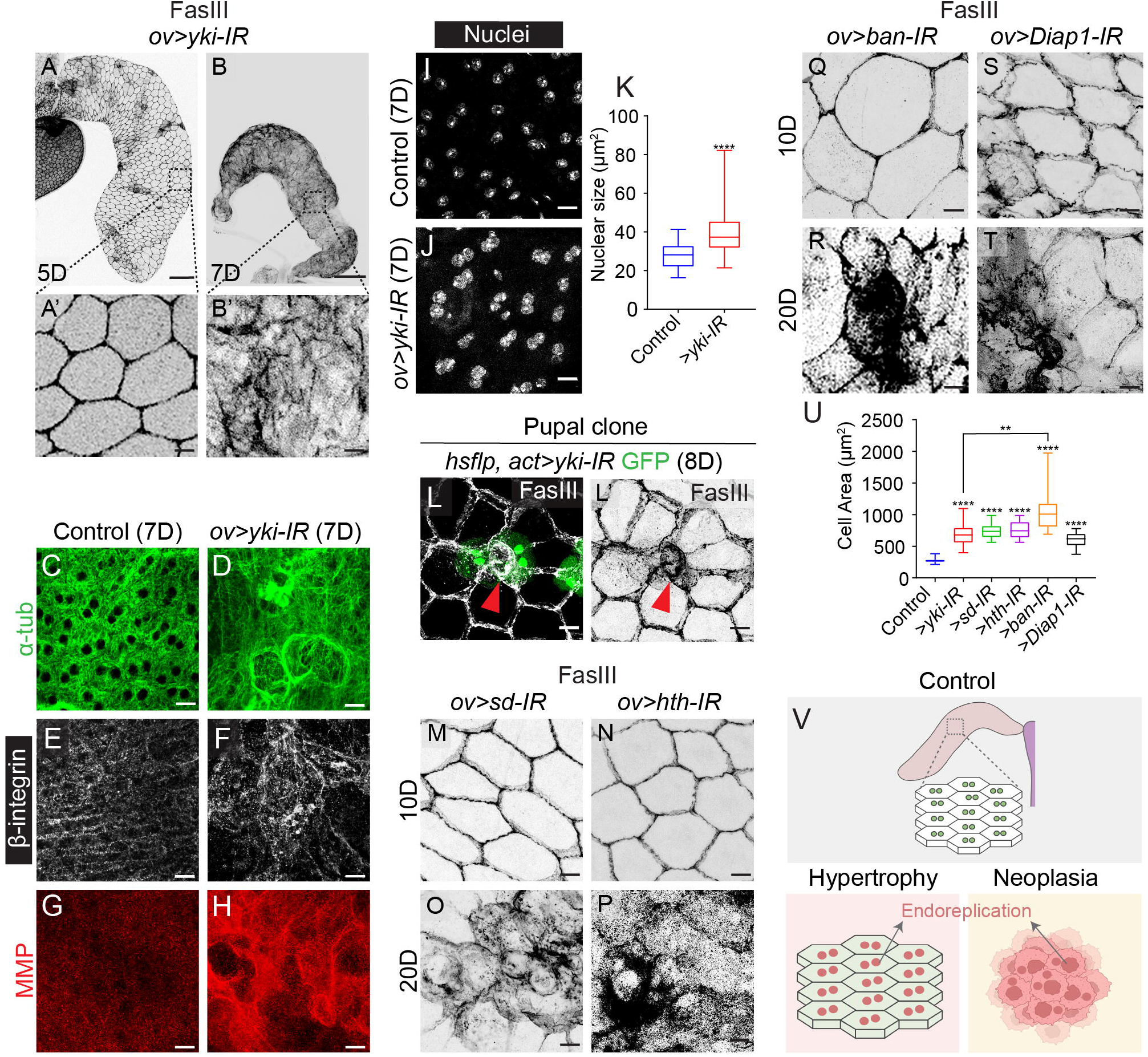
*yki* knockdown-induced cell hypertrophy and squamous cell carcinoma, SCC, in adult MAG. **(A-B)** Maximum intensity projections of optical sections of adult MAG stained for FasIII (grey) following *ov-Gal4*-induced knockdown of *yki* (*ov>yki-IR*) (A-B). Boxed areas are displayed at a higher magnification in the lower panel to reveal initial cell hypertrophy (A’) and, subsequently, SCC (B’), the latter displaying loss of the characteristic FasIII-marked ordered cell architecture. **(C-H)** Comparison of microtubule (C-D, green, α-tubulin), β-integrin (E-F, Mys, white), and matrix metalloprotease (G-H, MMP, red) in control and MAG SCC. **(I-K)** Comparison of nuclear size (TO-PRO, white) between control (I) and MAG SCC (J, *ov>yki-IR*) and their quantifications (K). Unpaired *t-*test of average nuclear size between control and *ov>yki-IR*, \*\*\*\**p*<0.0001; each data point represents mean ± *SD* (*n*= 50). **(L)** Somatic clonal (GFP, green) knockdown of *yki* in pupal columnar MAG *(hs-flp, act>yki-IR)* display rosette formation and delamination (red arrowhead). **(M-P)** FasIII-stained MAG displaying knockdown *sd* or *hth* display initial cell hypertrophy (10D, M-N) and, subsequently, SCC (20D, O-P). **(Q-T)** Knockdown of *ban* (Q, R) or *Diap1* (S, T) in adult MAG displayed initial cell hypertrophy and subsequent SCC. **(U)** Quantifying cell areas in MAG showing hypertrophy; unpaired *t*-test of average cell area between control and test genotypes \*\*\*\**p*<0.0001, while that between *yki* and *ban* knockdowns was \*\**p*<0.0016. Each data point represents mean ± *SD* (*n*=50). (**V)** Cartoon representation of *yki*-knockdown-induced early hypertrophy and, subsequently, SCC. Scale bar: (A-B) 100 µm, (A’, B’, C-T) 10 µm.

By comparison to Yki signaling loss-induced SCC (**Fig. 2**), its gain in adult MAG caused a reduction in both cell and organ sizes (*ov>yki^3SA^,* **Fig. S2**). Further, loss of Yki signaling in the columnar epithelium of pupal MAG resulted in progressive cell shrinkage and delamination (*hs-flp, act>yki-IR*; **Fig. 2L**, also see (Athilingam et al., 2022; Toyama et al., 2008). By contrast, it’s gain in pupal MAG columnar epithelium induced cell hypertrophy and neoplastic transformation (*hs-flp, act>yki^3SA^*; **Fig. S2**). These outcomes of loss or gain Yki signaling in pupal MAG are reminiscent of that seen in columnar epithelia of larval imaginal discs (Huang et al., 2005) or that seen in adult intestinal epithelium (Kwon et al., 2015; Staley and Irvine, 2010).

To further test if loss of Yki downstream targets (Nolo et al., 2006; Peng et al., 2009; Wu et al., 2008) induce MAG SCC, we first individually knocked down nuclear partners of Yki, like the TEAD domain, Scalloped, Sd (Goulev et al., 2008; Zhang et al., 2008) or TALE-homeodomain Homothorax (Hth) transcription factor (Peng et al., 2009). Knockdown of Sd (*ov>sd-IR*, **Fig. 2M**) or Hth (see *ov>hth-IR*, **Fig. 2N**) induced hypertrophy of MAG SE by seven days of adult life, which turned into SCC in 20-days-old adults (**Fig. 2O-P**). Thus, as in the peripodial epithelium of the wing imaginal disc (Friesen and Hariharan, 2023), Sd and Hth transcription factors are active in the MAG SE. A comparable—but slow-paced— hypertrophy-leading-to-SCC was also seen following the knockdown of other downstream targets of Yki, for instance, *bantam* (Brennecke et al., 2003) (*ov>ban-IR*, **Fig. 2Q-R**) or *Diap1* (Wu et al., 2003) (*ov>Diap1-IR,* **Fig. 2S-T**). We also noticed a more pronounced SE cell hypertrophy (**Fig. 2U**) and an increase in MAG organ sizes (**Fig. S2**) upon *ban* knockdown than those seen upon *yki, sd,* or *Diap1* knockdowns; these results appear consistent with a selective role of *ban* in the regulation of tissue growth (Brennecke et al., 2003; Nolo et al., 2006; Thompson and Cohen, 2006). In summary, compromising Yki signaling causes MAG SE cell hypertrophy, which culminates in SCC (**Fig. 2V**).

Does Yki-loss-induced MAG SCC qualify as true cancer? Like their human counterparts, a hallmark of lethal cancers in *Drosophila* is their ability to induce progressive loss of body mass, or cachexia, in the host adults (Figueroa-Clarevega and Bilder, 2015; Kwon et al., 2015). Crucially, cachexia-induced death of a patient or in *Drosophila* is not linked to cancer metastasis *per se* (for review, see (Argilés et al., 2018)). Adults displaying Yki loss-induced MAG SCC (*ov>yki-IR*) were marked by bloated abdomen (**Fig. 3A-B**), atrophy of abdominal muscle (**Fig. 3C-D**), and loss of fat body, characterized by decrease in lipid content (**Fig. 3E-F**): these are some of the telltale signatures of cancer-induced cachexia in the host animal (Figueroa-Clarevega and Bilder, 2015; Kwon et al., 2015), for review, see (Liu et al., 2022)). Previously, it was shown that tumor-derived Impl2, a *Drosophila* homolog of mammalian insulin-like growth factor binding proteins (IGFBPs, see (Andersen et al., 2000))—underpins cancer cachexia (Figueroa-Clarevega and Bilder, 2015; Kwon et al., 2015). In agreement, we noticed upregulation of ImpL2-GFP, a GFP fusion protein (Nagarkar-Jaiswal et al., 2015) (*ImpL2-GFP; ov> yki-IR;* **Fig. 3G-H**) in MAG-SCC. Finally, the knockdown of *ImpL2* in MAG SCC suppressed cachexia in the host adults (*ov>impL2-IR, yki-IR*, **Fig. 3I-J**)—despite their persistent cancerous growth (**Fig. 3K-K’**)—and with an accompanying restoration of their lifespan (**Fig. 3L**).

**Fig. 3.**
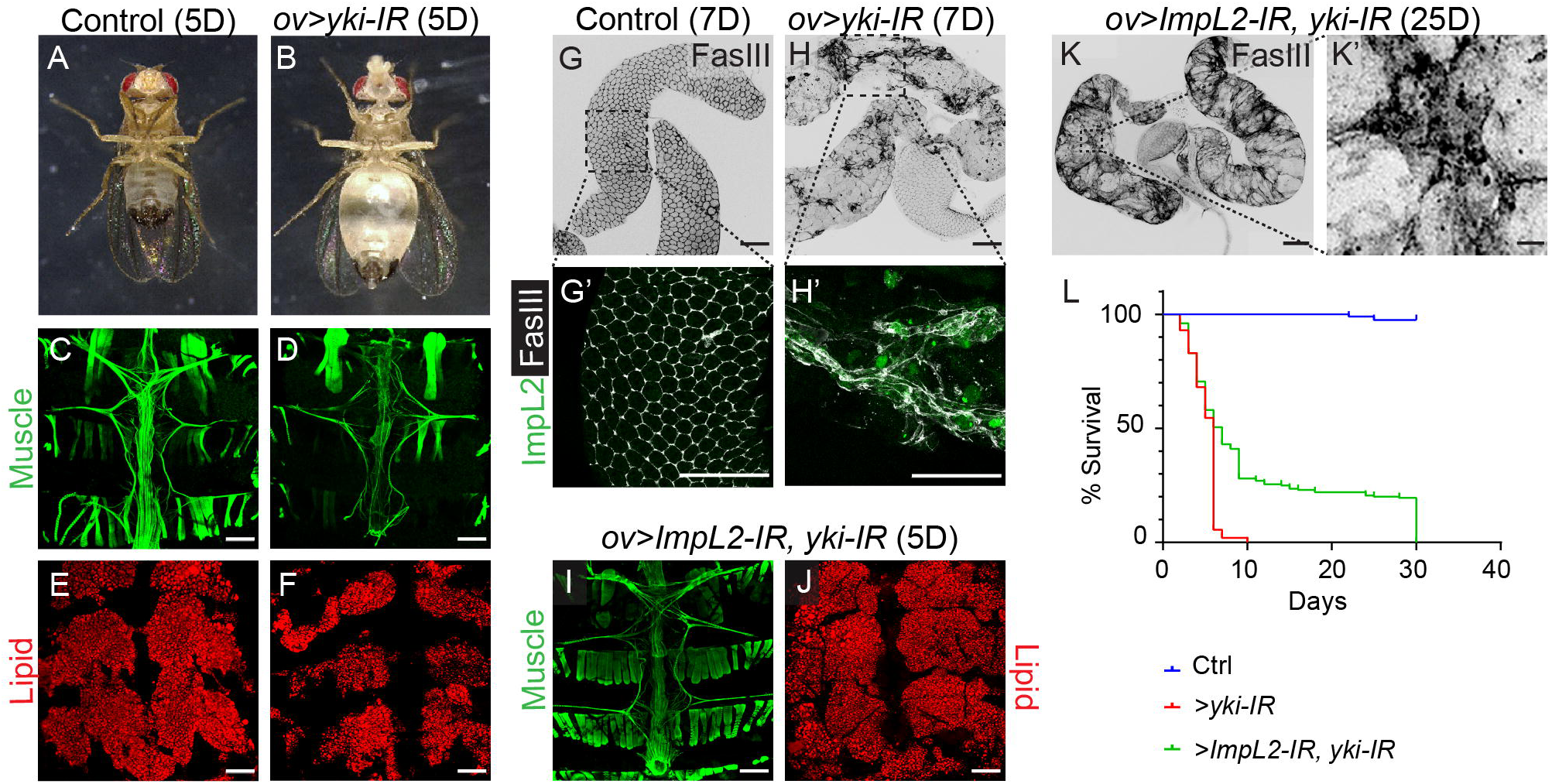
MAG SCC induced by loss *yki* signaling displays ImpL2-induced cachexia and host lethality. **(A-F)** Control (A) and test adults with *yki-*knockdown-induced MAG SCC(B); note the bloated abdomen in the latter. Respective abdominal muscles are stained for muscle (C-D, Phalloidin, green) and lipid (E-F, Nile red). (**G-H**) Comparison of ImpL2-GFP (green) expression in control MAG (G) and *yki*-knockdown-induced MAG SCC (H). Boxed areas from these MAGs are shown at higher magnification in the respective bottom panels (G’-H’). **(I-K)** Knockdown of *ImpL2* from *yki-*knockdown-induced MAG SCC reversed the atrophy of host adult abdominal muscle (green) (I), besides restoring lipid content (red) in its fat body (J), but without arresting SCC (FasIII, K) *per se*. The boxed area in (K) is shown at high magnification in (K’). (**L**) Kaplan-Meier survival curves of these animals or upon knockdown of *ImpL2*. Scale bar: (A-K, L) 100 µm, (K’, L’)10 µm.

Together, these results reveal that Yki is a tumor suppressor in *Drosophila* adult MAG and that loss of its signaling causes lethal, cachexia-inducing SCC.

### Gain of oncogenic Ras^V12^ signaling in the SE of adult MAG does not induce SCC

Basal extrusion of oncogenic Ras^V12^ expressing MAG cells was recently modeled in *Drosophila* as a precursor of their cancerous transformation and reminiscent of human prostate cancer (Rambur et al., 2020). In that study, the authors induced gain of Ras^V12^ in the pupal MAG (see Fig. 2D-E, (Rambur et al., 2020)) when, as shown here, cells are columnar (see **Fig. 1B, Fig. S1**). We could recapitulate these results in the adult MAG by inducing clonal initiation of Ras^V12^ signaling in the pupal columnar epithelium of MAG. Such Ras^V12^-expressing clones displayed altered cytoarchitecture, revealing their cancerous transformation (**Fig. S3**). By contrast, induction of Ras^V12^ under the *ov-Gal4* driver (*ov>Ras^V12^*)—that is, following cell-flattening-induced nuclear Yki signaling in the MAG epithelium—failed to induce neoplasia, except mild hypertrophy (**Fig. 4A-D**) and rosette-like cell arrangements at later stages (**Fig. 4E-F**), an indication of delamination due to cell death (Athilingam et al., 2022; Toyama et al., 2008). Cell (**Fig. 4G**) and organ hypertrophy (**Fig. 4H**) in the Ras^V12^ expressing adult MAGs appear generally insignificant (**Fig. 4A-D, Fig. S3**) except after prolonged expression in older adults (**Fig. 4E-F**; **Fig. S3**). Crucially, adult hosts bearing Ras^V12^-expressing MAGs did not display cachexia (**Fig. 4I-J’**) or shortening of lifespan (**Fig. 4K**), unlike their counterparts exhibiting knockdown of Yki signaling (see **Fig. 3**). These outcomes of Ras^V12^ expression in adult MAGs also appeared consistent with their persistent nuclear Yki signaling as seen from their *Diap1-lacZ* reporter expression (*Diap1-lacZ; ov>Ras^V12^*, **Fig. 4L-O**). Surprisingly, Ras^V12^-expressing adult MAG displayed enlarged nuclear volume, suggesting their endoreplication (**Fig. 4P**), reminiscent of those showing knockdown of *yki* (see **Fig. 2J-K**), revealing thereby that endoreplication without accompanying cytoarchitectural changes does not constitute MAG carcinogenesis.

**Fig. 4.**
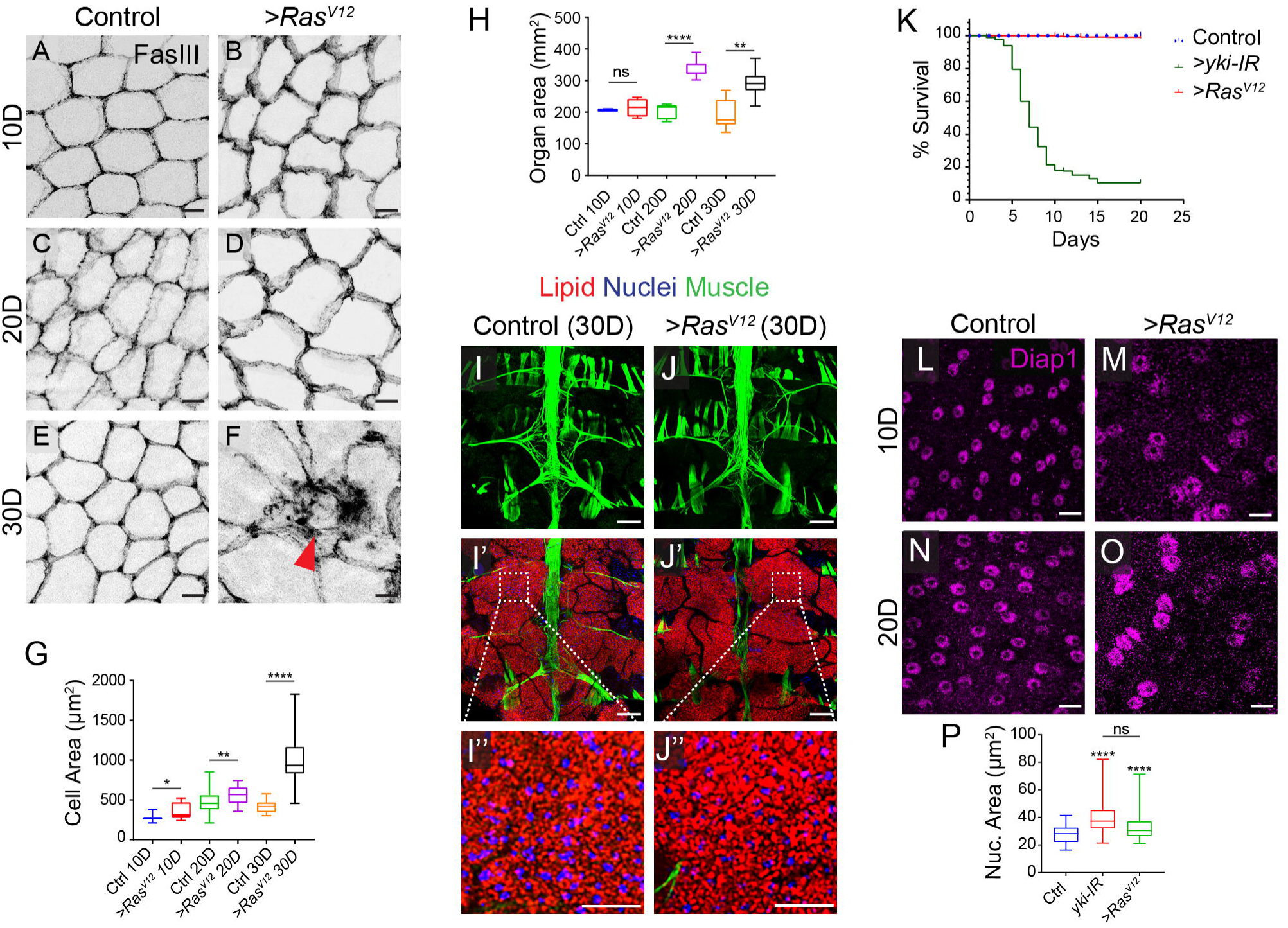
Oncogenic Ras^V12^ does not transform the squamous cells of adult MAG. **(A-F)** Control and *ov-Gal4>Ras^V12^* MAG epithelium from adults 10D (A, B), 20D (C, D), or 30D (E, F) days after eclosion. **(G-H)** Quantification of their cell areas (G) and analysis by unpaired *t*-test from 10D (\**p*=0.0170), 20D (\*\**p*=0.0011), and 30D (\*\*\*\**p*<0.0001) samples. Likewise, the comparision of organ sizes at 10D (*ns*, *p*=0.4345), 20D (\*\*\*\**p*<0.0001), and 30D (\*\**p*=0.0013) (H). Note the rosette-like appearance and cell delamination (red arrowhead) in 30D-old Ras^V12^ expressing MAG. **(I-J)** Abdominal muscle (actin, green) and fat body (stained for Nile red and nuclei, TO-PRO, blue) of control (I) and *ov-Gal4>Ras^V12^* (J) MAG-bearing 30 days old adults. Boxed areas in the middle panel (I’-J’) are shown at higher magnification at the bottom (I’’-J”). **(K)** Kaplan-Meier survival curves of adults with control (blue), *ov-Gal4>yki-IR* (green), and *ov-Gal4> Ras^V12^* (red) MAG. **(L-O)** *Diap1-lacZ* (β-gal, magenta) expression in control and *ov-Gal4>Ras^V12^* MAG from 10D (L-M) and 20-day-old (N-O) adults. **(P)** Quantification of the nuclear area of indicated genotype (*n*=50) by unpaired *t*-test between control and *ov>yki-IR* (\*\*\*\**p*<0.0001), between control and *ov>Ras^V12^* (\*\*\*\**p*<0.0001), and between *ov>yki-IR* and Ras^V12^ (*p*=0.0547); the latter was not significant (ns). Scale bar: (A-F) 10 µm, (I-J’’) 100 µm, (L-O) 10 µm.

Thus, contrary to the catastrophic fallout of knockdown of Yki signaling, a gain of Ras^V12^ in the SE of adult MAG is not oncogenic. Therefore, the previous report on Ras^V12^-induced MAG carcinogenesis (Rambur et al., 2020) essentially represents the transformation of its pupal columnar epithelium.

### Yki loss-triggered PI3K/Akt/TOR signaling induces MAG cell hypertrophy and SCC

Nutrient-sensitive PI3K/Akt/TOR pathway is an evolutionarily conserved cell and organ size regulator (for review, see (Condon and Sabatini, 2019; Wullschleger et al., 2006)). MAG epithelial cells are nutrient-sensitive (Kubo et al., 2018). Not surprisingly, starved adult flies showed a decrease in cell sizes in the MAG SE compared to their fed counterparts (**Fig. 5A-B’**). Given that Yki signaling-compromised MAG display cell hypertrophy preceding their progression to SCC (**Fig. 2**), we further reasoned recruitment of TOR signaling (for review, see (Condon and Sabatini, 2019)) during MAG SE transformation. Indeed, crosstalks between Yki/YAP/TAZ and TOR signaling pathways have previously been shown to regulate organ size in most epithelial organs (Parker and Struhl, 2015; Tumaneng et al., 2012a; Tumaneng et al., 2012b). We thus first tested the consequence of gaining or knocking down the TOR signaling pathway in the adult MAG. We noticed that the knockdown of PTEN, a negative regulator of TOR (*ov>PTEN-IR*, **Fig. 5C**, **Fig. S4**), increased cell sizes, as can be expected from enhanced TOR signaling in these cells. By contrast, the knockdown of phosphoinositide-3-kinase (*ov>PI3K-IR,* **Fig. 5D**, **Fig. S4**), the upstream positive regulator of the TOR signaling pathway for cell growth (Zhang et al., 2000)) or TOR (*ov>TOR^DN^*, **Fig. 5E**)—each individually compromising TOR signaling—decreased MAG SE cell sizes (**Fig. 5F**).

**Fig. 5.**
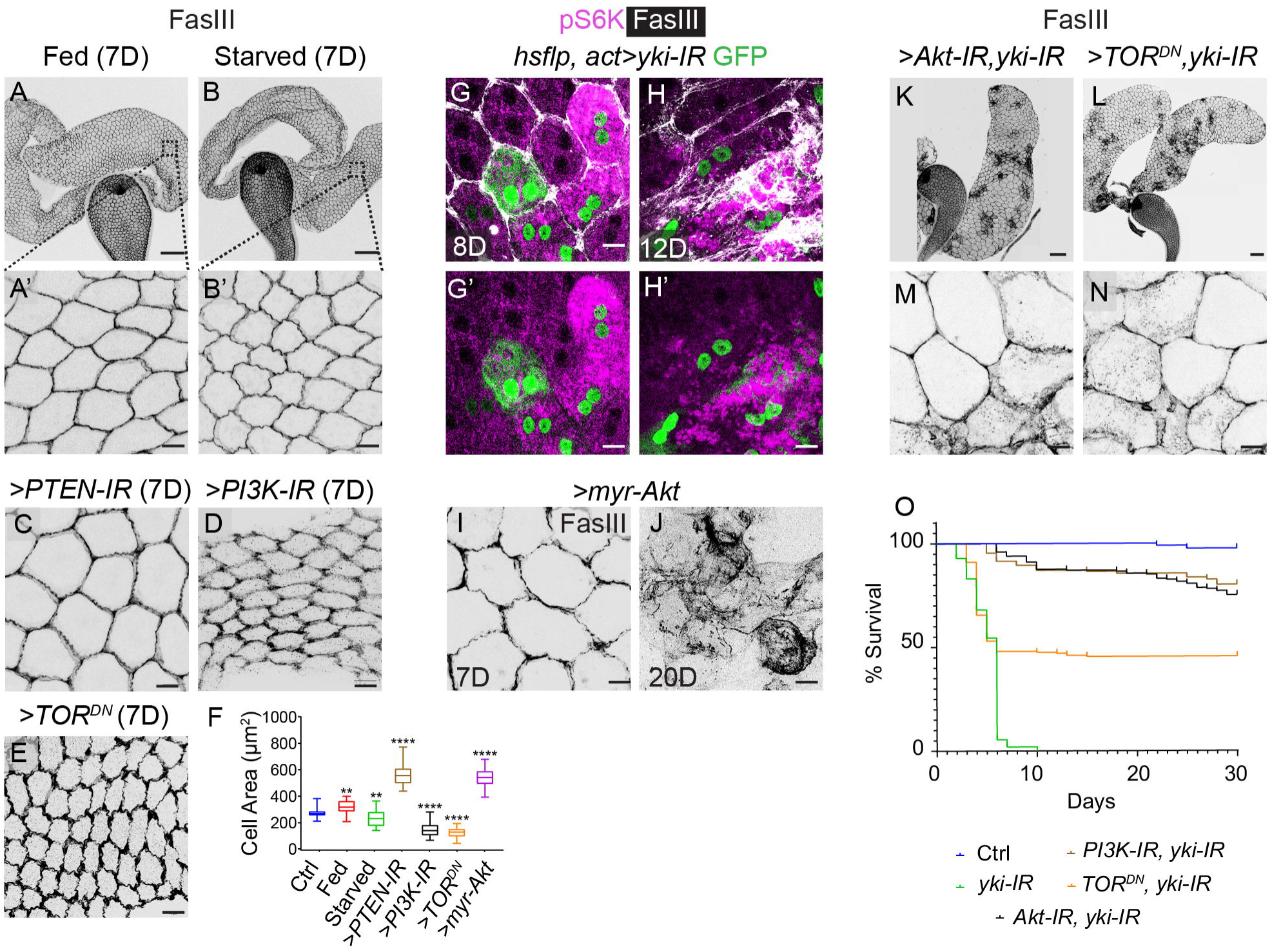
Knockdown of *yki* triggers TOR-induced adult MAG SCC. **(A-B)** Whole mounts of MAG, immunostained for FasIII (grey), from fed (A) and starved adults (B). The lower panel (A’-B’) displays representative magnified areas for comparison of cell sizes. **(C-F)** Comparison of cell areas of 7 days old adult MAG from *ov-Gal4>PTEN-IR* (C), ov*-Gal4>PI3K-IR* (D), and *ov-Gal4>TOR^DN^* (E) and their quantification (F). Unpaired *t*-test of average cell area (*n*=50) between control and fed (\*\**p*=0.0011), between control and starved (\*\**p*=0.0011), and between control and rest of the indicated genotypes (\*\*\*\**p*<0.0001). **(G-H)** GFP-marked somatic clones expressing the *yki-IR* transgene, induced in 5-day-old adult MAG and dissected at 8D (G, G’), revealed cell hypertrophy and upregulated pS6K (magenta). In contrast, those examined at 12D (H, H’) display and SCC (H, FasIII, white) upregulated pS6K apart. (**I-J**) *ov-Gal4>myr-Akt* MAG display initial hypertrophy (I, 7D) and, subsequently, SCC (J, FasIII, 20D). (**K-N)** *ov*>*Akt-IR, yki-IR* (K), and *ov*> *TOR^DN^, yki-IR* MAG (L) do not display SCC and are instead marked by cell hypertrophy (M, N). **(O)** Kaplan-Meier survival plot of the indicated genotypes. Scale bars: (A-B) 100 µm, (A’-J) 10 µm, (K, L) 100 µm, (M, N) 10 µm.

We also noticed that somatic clones displaying knockdown of Yki signaling in adult MAG SE (*hs-flp, act>yki-IR*)—and thereby its cell hypertrophy and SCC—showed elevated pS6K (Miron et al., 2023), a TOR target, **Fig. 5G-H’**). Further, expression of a constitutively active form of Akt, myr-Akt—that elevates TOR signaling (Stocker et al., 2002)—induced hypertrophy of MAG SE (*ov>myr-Akt*, **Fig. 5I, Fig. S4)** and eventually MAG SCC **(Fig. 5J**). Finally, compromising TOR signaling via knockdown of PI3K (*ov>PI3K-IR, yki-IR*, **Fig. S4**), Akt (*ov>Akt-IR*, *yki-IR*, **Fig. 5K**) or TOR (*ov>TOR^DN^, yki-IR*, **Fig. 5L**) arrested MAG SCC (**Fig. 5M-N**), and improved adult lifespan (**Fig. 5O**).

Thus, an upregulation of the PI3K/Akt/TOR pathway is causally linked to Yki loss-induced MAG SE hypertrophy and SCC.

### Yki signaling maintains squamous cell homeostasis in larval tracheal tubes and Malpighian tubules

The fallouts of loss of Yki signaling in the MAG SE, namely, cell hypertrophy and SCC (**Fig. 2–4**), are surprisingly different from that reported in the follicular epithelium of the egg chambers (Fletcher et al., 2018) or peripodial membrane reported previously (Fletcher et al., 2018; Friesen and Hariharan, 2023). Although MAG SE displays nuclear Yki signaling, like those of its follicular or peripodial counterparts, the anatomical context of their cell flattening and mechanical strains registered in the lining of the lumen in a tubular organ are likely different. This scenario appears reminiscent of disparate outcomes of loss of YAP/TAZ in a diverse type of flattened and mechanostrained human cells. For instance, YAP/TAZ loss-induced lung SCC (Huang et al., 2017), alveolar cysts in mammalian lungs (Mahoney et al., 2014; van Soldt et al., 2019), loss of differentiation of flattened retinal pigment epithelium (Kim et al., 2016), or arrested vasculogenesis when lost in blood vessel endothelial cells (Ong et al., 2022). Therefore, we asked if SEs lining the lumens of other tubular organs in *Drosophila* are susceptible to loss of Yki signaling, like that of MAG. We thus knocked down Yki signaling in the SE linings of other tubular organs in *Drosophila,* like the larva tracheal tubes (Affolter and Shilo, 2000; Ghabrial et al., 2003; Metzger and Krasnow, 1999) or adult Malpighian tubules (Jung et al., 2005).

A pair of major tracheal tubes, termed dorsal trunks (DT), run through the entire length of the larvae—anterior-to-posterior—sending out lateral and dorsal branches that further ramify into increasingly narrower units to finally culminate in tracheoles, which deliver oxygen to the receiving cells through diffusion (Hayashi and Kondo, 2018; Sato and Kornberg, 2002; Scholl et al., 2021). The SE lining the air-filled lumen of the DT displays nuclear translocation of Yki (**Fig. 6A-B”**), marked by nuclear *ban-lacZ* (**Fig. 6C**) and *Diap1-lacZ* (**Fig. 6D**), coinciding with their cell flattening during the early third instar larva— transitioning from its cytoplasmic localization in the preceding second larval instar (Yki::GFP, **Fig. 6A-B”, S5**). We used a *breathless*, *btl-Gal4* driver (Shiga et al., 1996) to selectively knockdown Yki signaling following its nuclear translocation in the early third larval instar (**Fig. S5**). We note that in comparison to control (**Fig. 6E**), loss of Yki signaling (*btl-Gal4, Gal80^ts^>yki-IR*, **Fig. 6F**), or its downstream target, *ban* (*btl-Gal4, Gal80^ts^>ban-IR*, **Fig. 6G**) in the SE of DT resulted in cell hypertrophy and loss of FasIII localization. However, hypoxia was not seen in *yki* or *ban* knockdown in third instar larvae compared to control (**Fig. S5**). Reminiscent of MAG-SCC rescue, the knockdown of TOR signaling reversed Yki loss-induced DT cell hypertrophy (**Fig. 6H**). We also notice an increase in nuclear size and fluorescence intensity (**Fig. 6I-K**), a hallmark of their cancerous transformations (see Fig. 3 (Levine and Cagan, 2016), also see (Alonso-Lecue et al., 2017; Zhang et al., 2022)), that gets restored upon downregulation of TOR signaling (**Fig. 6L-N**).

**Fig. 6.**
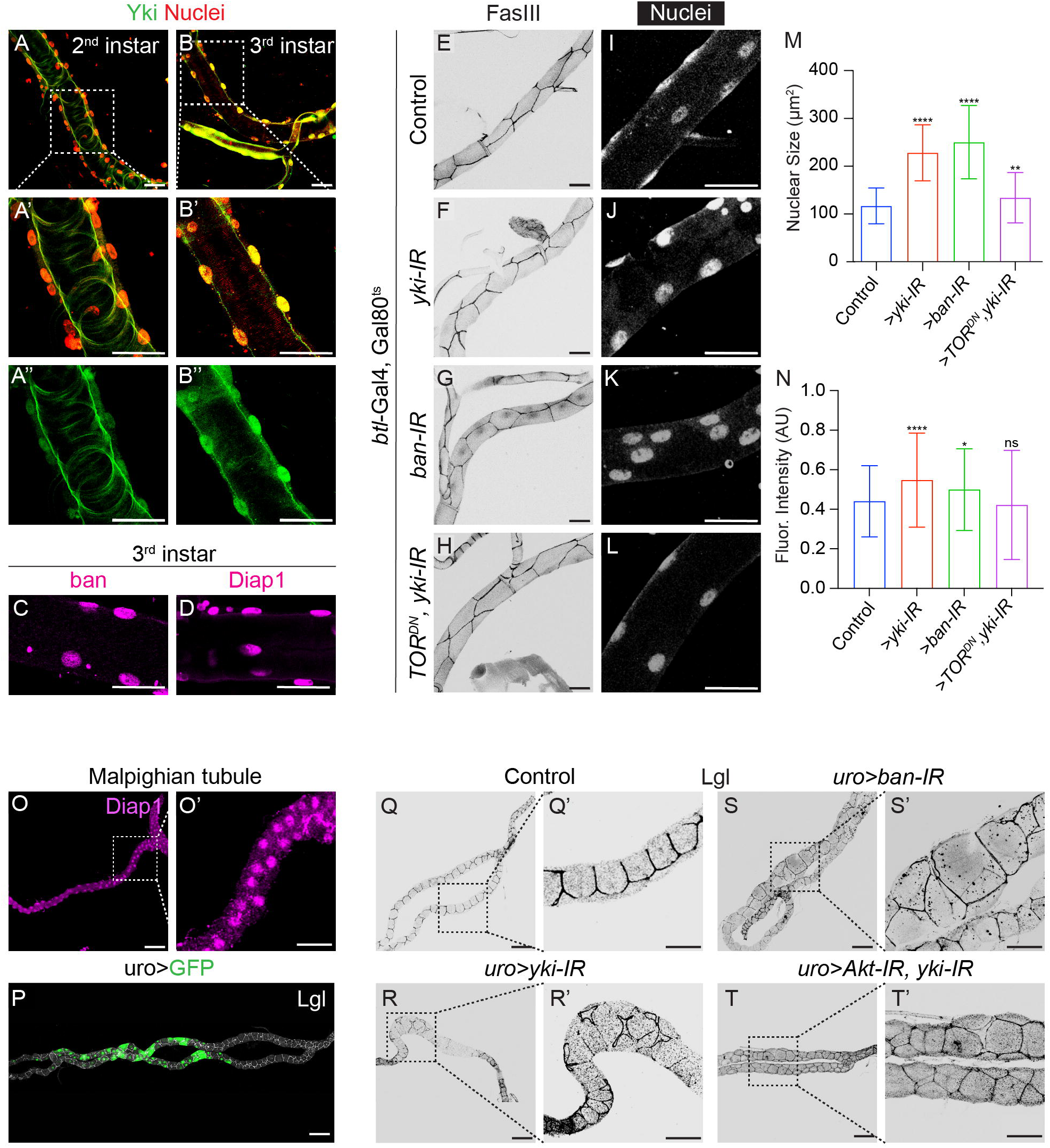
Yki signaling maintains squamous cell homeostasis in larval tracheal tubes and Malpighian tubules. (**A-B**) Yki::GFP (green) in the dorsal tracheal trunk (DT) from the second (A) and third instar larva (B); nuclei are marked by TO-PRO (red). Boxed areas from the merged image are shown at higher magnification in the panels below (A’-B’) to reveal cytoplasmic (A”) and nuclear localization of Yki::GFP (B”) in the DT from the second and third instar larva, respectively. (**C-D**) Yki signaling targets, *ban* (C, *ban-lacZ,* magenta), and *Diap1* (D, *Diap1-lacZ,* magenta) displaying nuclear localization in the DT of third instar larvae. (**E-L**) Third instar larval tracheal DT of control (E), *btl-*Gal4, Gal80^ts^>*yki-IR* (F), *btl-*Gal4, Gal80^ts^ *>ban-IR* (G) and *btl-*Gal4, Gal80^ts^ *>TOR^DN^, yki-IR* (H) stained for FasIII (grey) and nucleus (I-L, TO-PRO, white). **(M-N)** Quantification of nuclear size (M) and fluorescence intensity (N) of indicated genotypes. Unpaired *t*-test of average nuclear size (*n*=50) between control and *yki* loss (\*\*\*\**p*<0.0001), between control and *ban* loss (\*\*\*\**p*<0.0011), between control and *btl-*Gal4, Gal80^ts^ *>TOR^DN^, yki-IR* (**p<0.0045). **(O)** Malpighian tubules from adults displaying nuclear localization *Diap1-lacZ* (β-gal, magenta). The boxed area in this figure is magnified on the right panel (O’). **(P)** *uro-Gal4* expressing (GFP) domain in Malpighian tubules. **(Q-T)** Malphigian tubules of control (Q), *uro>yki-IR* (R), *uro>ban-IR* (S), and *uro>Akt-IR, yki-IR* (T) immunostained for Lgl (grey). Box areas in these images are shown at higher magnification in respective right panels (Q’-T’) to reveal cell hypertrophy in the Malpighian tubule upon knockdown of *yki* (R’) or its target *bantam* (S’) and suppression of their cell hypertrophy upon simultaneous knockdown of TOR signaling (T’). Note the loss of Lgl from the cell boundaries in the *uro>yki-IR* Malpighian tubule, indicating neoplasia. Scale bars: (A-L) 50 µm, (O-T) 100 µm and insets 50 µm.

Like MAG, the SE-lined lumen of *Drosophila* adult Malpighian tubules (**Fig. S5**) displays perpetual Yki signaling, marked by nuclear *Diap1-lacZ* expression (**Fig. 6O-O’**). Knockdown of *yki* or *ban* under the *uro-Gal4* driver—that expresses in the main segment of Malpighian tubules (**Fig. 6P**; see (Terhzaz et al., 2010))—induced cell hypertrophy and loss of a basolateral protein and a tumor suppressor, Lgl (Bilder et al., 2000) (**Fig. 6Q-S’**). We also notice an increase in nuclear size (**Fig. S5**) and elevated cytoplasmic phosphorylated 4EBP levels (**Fig. S5**) upon *ban* knockdown (*uro>ban-IR*). Finally, the loss of Akt rescues *yki* loss-induced hypertrophy in the Malpighian tubule (**Fig. 6T-T’**).

These findings, therefore, reveal a pervasive role of nuclear *yki* as a tumor suppressor of squamous epithelia in multiple tubular organs in *Drosophila*.

## DISCUSSION

Ascertaining the instructive role of a given signaling pathway demands a critical examination of its genetic loss. A careful appraisal of the loss of highly conserved YAP/TAZ/Yki signaling did not support the notion of their endogenous regulation of organ growth (Kowalczyk et al., 2022). Instead, its developmentally regulated perpetual activations (Fletcher et al., 2018; Friesen and Hariharan, 2023; Kumar et al., 2019; Mahoney et al., 2014; Moreno-Marmol et al., 2018) likely regulate the homeostasis of mechanostrained SE in diverse organs. Here, we have tested this possibility and showed that endogenous Yki signaling negatively regulates the PI3K/Akt/TOR signaling, thereby maintaining cell and organ homeostasis in the SE-lined lumens of three tubular organs of *Drosophila*. Thus, genetic knockdown Yki signaling triggers elevated PI3K/Akt/TOR signaling—as in the adult MAG SE—inducing cell hypertrophy, culminating in SCC. Thus, while an out-of-context gain of Yki signaling often turns cancer-promoting in organs with non-squamous epithelium (Harvey et al., 2013; Song et al., 2019; Tumaneng et al., 2012a; Zanconato et al., 2016), it is a knockdown of Yki signaling as revealed here—and reminiscent of loss of a tumor suppressor—which turns oncogenic in the SE-lined lumens of tubular organs of *Drosophila*. Remarkably, host adults bearing MAG SCC display extreme cancer cachexia (Ding et al., 2021; Fearon et al., 2012; Figueroa-Clarevega and Bilder, 2015; Kwon et al., 2015; Song et al., 2019) marked by their early mortality. Thus, Yki-compromised MAG SCC likely represents one of the most lethal cancers in *Drosophila*. Notably, our results also represent the first instance of a genetic modeling of SCC in *Drosophila*.

Strikingly, our results reveal that cells from a single developmental lineage may display cell shape/architecture-dependent fallout of the knockdown of Yki signaling. Thus, while the gain of Yki signaling in the columnar pupal MAG is oncogenic, its loss induces SCC in their adult squamous counterparts. This disparate, cell shape-/architecture-dependent response to potentially oncogenic signaling is further underscored by the observation that oncogenic Ras^V12^ signaling, too, induces cancerous transformation in the columnar pupal MAG epithelium but not in their squamous adult counterpart. It was shown previously that in non-tubular organs like adult follicular or larval peripodial SE (Fletcher et al., 2018; Friesen and Hariharan, 2023)—knockdown of Yki signaling do not induce cancerous transformation, unlike that seen in the SE of tubular MAG, which further underscore the importance of the tissue architecture-dependent roles of Yki signaling. Additionally, even amongst the tubular organs tested here, the entire gamut of fallouts of loss of Yki signaling in the SEs are not identical. Thus, while loss of epithelial junctional architecture, cell hypertrophy, and endoreplication are shared features of knockdown of Yki signaling in all three tubular organs examined here, cachexia-inducing lethal SCC is seen in only the adult MAG.

We further reveal that crosstalks between two highly conserved signaling pathways regulating organ size—Hippo and PI3K/Akt/TOR (Tumaneng et al., 2012a)—drive MAG SCC when deregulated. PI3K/Akt/TOR signaling activating upon knockdown of *yki* is likely the predominant causal underpinning of MAG SCC. Suppression of MAG SCC upon knockdown of PI3K/Akt/TOR signaling in Yki signaling-deficient MAG and, alternatively, PI3K/Akt/TOR-induced hypertrophy-culminating-in-MAG-SCC provides two complementary evidences to support this mechanism. Crosstalks between Hippo and PI3K/Akt/TOR signaling have been reported in multiple contexts (for review, see (Honda et al., 2023)), although unraveling their mechanistic underpinning continues to be a work in progress (Tumaneng et al., 2012b) (for review, see (Honda et al., 2023; Tumaneng et al., 2012a). For instance, transcriptional upregulation of YAP target microRNAs of the miR-29 family suppresses PTEN by targeting its 3’ UTR, activating TOR signaling in columnar epithelial cells (Tumaneng et al., 2012b). Previously, it has been shown that enhancement of PI3K/Akt/TOR and YAP/TAZ signaling takes place when cells are grown on substrates with high-density collagen (Shea et al., 2018), suggesting their select tissue-substrate-linked crosstalks. In this regard, it is well-recognized that the nature of extracellular signals may trigger specific responses within cells (López-Maury et al., 2008): for instance, diverse gene expression programs based on disparate mechanosensory cues arising from substrate. The contrasting outcomes of *yki* knockdown in the SEs from non-tubular (Fletcher et al., 2018; Friesen and Hariharan, 2023) and tubular organs (this report) likely exemplify this scenario. Thus, from amongst the plethora of possible routes to activation of PI3K/Akt/TOR signaling (Huang and Fingar, 2014; Shimobayashi and Hall, 2014), we speculate that cytoarchitectural alterations upon loss of Yki signaling (Halder et al., 2012), as proximate causal underpinnings of induction of MAG hypertrophy, which culminate in SCC. Our genetic modeling of MAG SCC via gain of PI3K/Akt/TOR signaling sets the stage for deconstructing the complex network of cellular events downstream of Yki in the SEs.

Finally, Yki-loss-induced MAG SCC also means that pharmacological activators of YAP/TAZ could suppress select cancer types. For instance, Digitoxin—a cardiac glycoside commonly prescribed for treating congestive heart failure (Arispe et al., 2008), has also been shown to activate YAP for treating YAP-loss-driven lung SCC (Huang et al., 2017). An extension of our study would imply that inhibitors of YAP (Oku et al., 2015; Yong et al., 2021; Zanconato et al., 2016) may aggravate SCC in select SEs, thereby calling for cautions in their therapeutic exploration (see (Luo et al., 2023)).

## MATERIALS AND METHODS

### *Drosophila* maintenance and generation of somatic clones

*Drosophila* were maintained at 25 ± 1°C on standard food. GFP-labelled mitotic clones were generated using heat shock flp/FRT-flip-out technique (Xu and Rubin, 1993) in pupal and adult males. To generate somatic clones in pupal MAG, heat shock was given for 10 min at 50h APF (<u>A</u>fter <u>P</u>upa <u>F</u>ormation)(Rambur et al., 2020). For adult studies, five-day-old eclosed adult males were given heat shock for 5 min, and MAG were, dissected 8-10 or 12-20 days later for assessment of phenotypes after initial and extended period of clonal growth, respectively.

### Strategies for gene overexpression and knockdown studies

For *ov-Gal4* (Heifetz et al., 2000) and *uro-Gal4* (Terhzaz et al., 2010) driven gene overexpression or knockdown, five days old adult flies of relevant genotypes were shifted from 25 ± 1°C to 29 ± 1°C, which enhances Gal4 activity (Duffy, 2002). The time of examination of the phenotypes varried in different crosses and these details are mentioned in respective outcomes. For all larval trachea-related experiments, freshly collected eggs were allowed to eclosed at 25°C. For driving *yki-IR* or other transgenes under the *btl-Gal4, Gal80^ts^* drivers first instar larvae were cultured at 18°C till the end of second instar larva and then shifted to 29°C (Duffy, 2002) until their dissection at late third instar larval stage.

### Immunostaining, microscopy, quantification, and statistical analysis

Tissues were dissected in 1x Phosphate Buffer Saline (PBS), and fixed in 4% paraformaldehyde in 1x PBS. Tissues were permeabilized in PBST (1XPBS containing 0.02 % Triton-X-100). After blocking in 5% BSA for 2 hrs, these were incubated in primary antibody overnight at 4^0^C, washed three time in for 10 min each in PBST and, finally, incubated with appropriate fluorescent-tagged secondary antibodies (1:250, Invitrogen) for two hours at room temperature. The samples were washed three time in for 10 min each in PBST, followed by two washes for 10 min each in PBS, and counterstained with TO-PRO-3 and/or Alexa Fluor™ 555 Phalloidin (A34055, Invitrogen), wherever appropriate. Finally, the tissues were washed three time in for 10 min each in PBS, mounted in Vectashield®, and sealed with transparent nail-paint. Samples were imaged with a 20x or 63x oil immersion objective in a Leica SP5 laser scanning confocal microscope. Images were analyzed and processed using Leica confocal software LAS AF and FIJI.

Adult fly bright field imaging was done using Leica M205 FA stereomicroscope. All images were assembled using Adobe Photoshop CS-6.

Cell area and fluorescence intensity were calculated using FIJI. Freehand drawing tool was selected to trace the cell, organ, or nuclear boundaries and calculate its surface area and Fluorescence intensity, wherever necessary. A 2D Stitch plugin was used in FIJI to stitch the entire MAG organ maximum intensity projection images. All statistical analyses were performed in GraphPad 9.0. Unpaired *t*-tests were performed for quantification of cell, organ, and nuclear sizes. A cutoff of *p*<0.05 was used to define statistical significance in all graphical plots. Illustrations were made using BioRender.

### Adult fat body dissection and their phalloidin, Nile Red and nuclear staining

The adult abdominal dorsal cuticles were carefully removed in 1X PBS buffer using micro-scissors, avoiding detachment of the underlying adipose tissue (fat body). These cutcle with attached fat body were fixed with chilled 4% paraformaldehyde in PBS for 20 min and then washed twice in PBS. Fixed tissues were stained using solution of 0.005% (1:200) Nile Red 555 (Sigma; 19123) for 1 hour at room temperature. The samples washed three time in for 10 min each in PBS. Separately, such tissues samples were stained with Phalloidin and TO-PRO-3 for 1 hour each, at room temperature. The samples were mounted after washing washed three time in for 10 min each PBS and imaged immediately.

## KEY RESOURCE TABLE

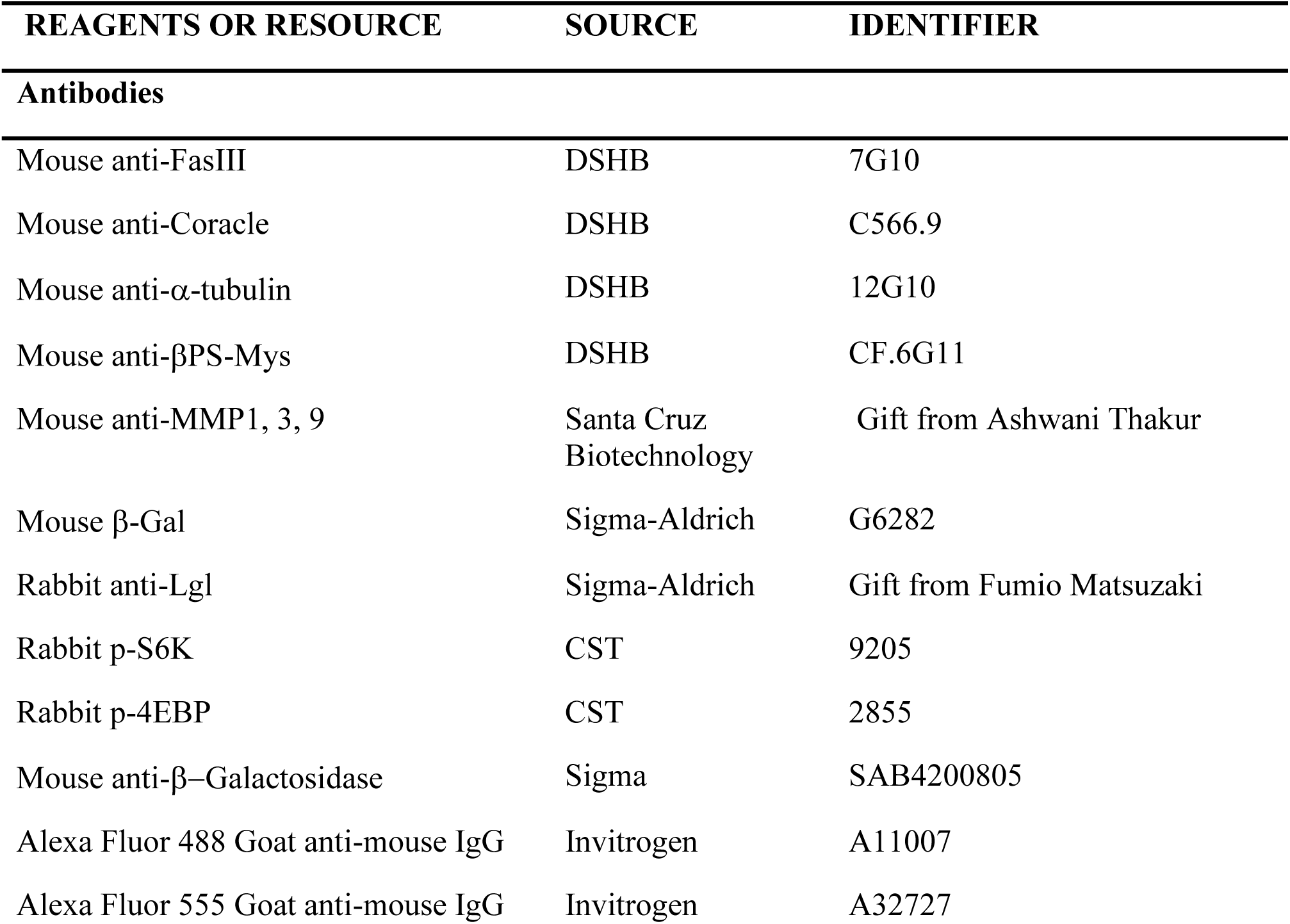
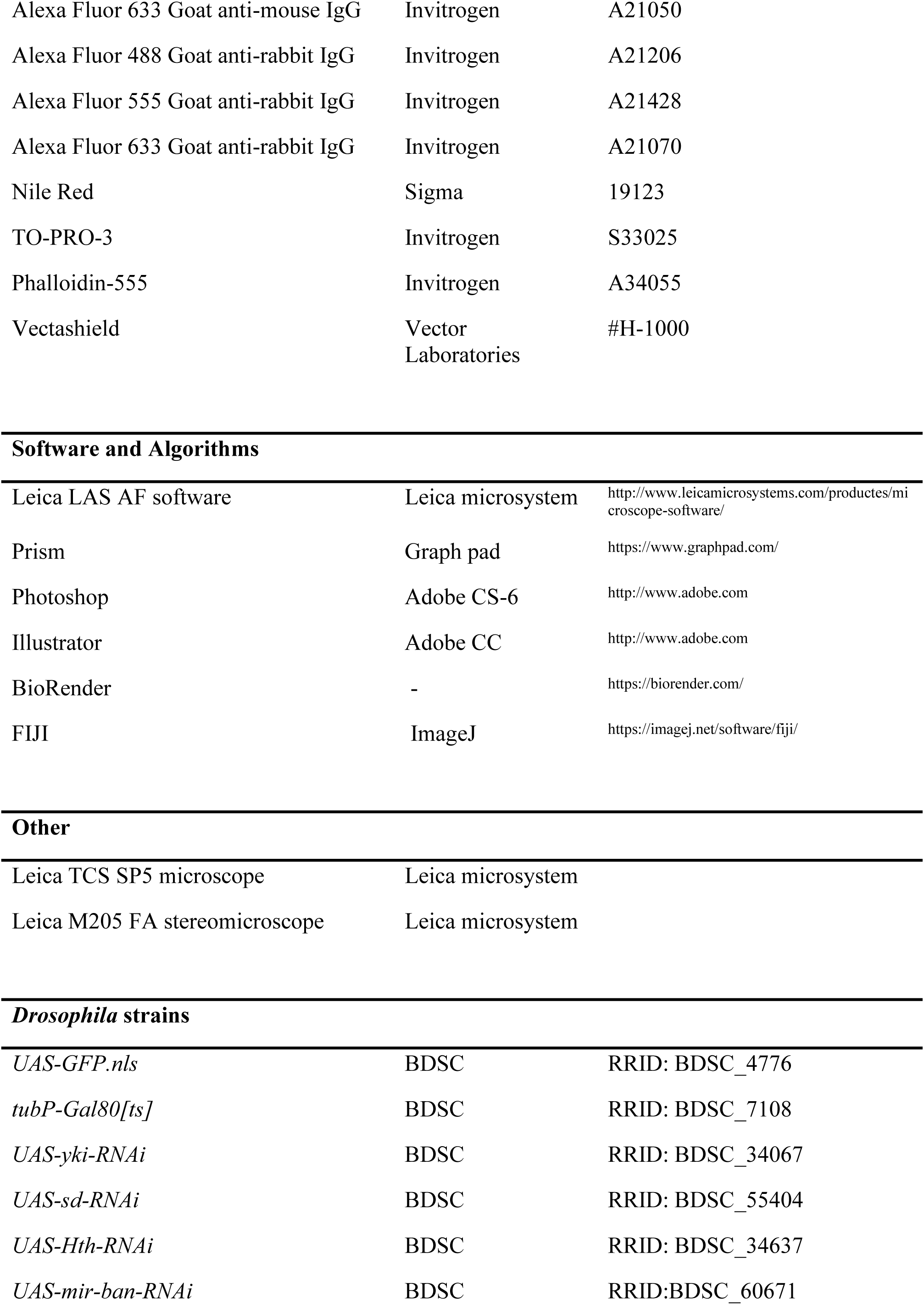
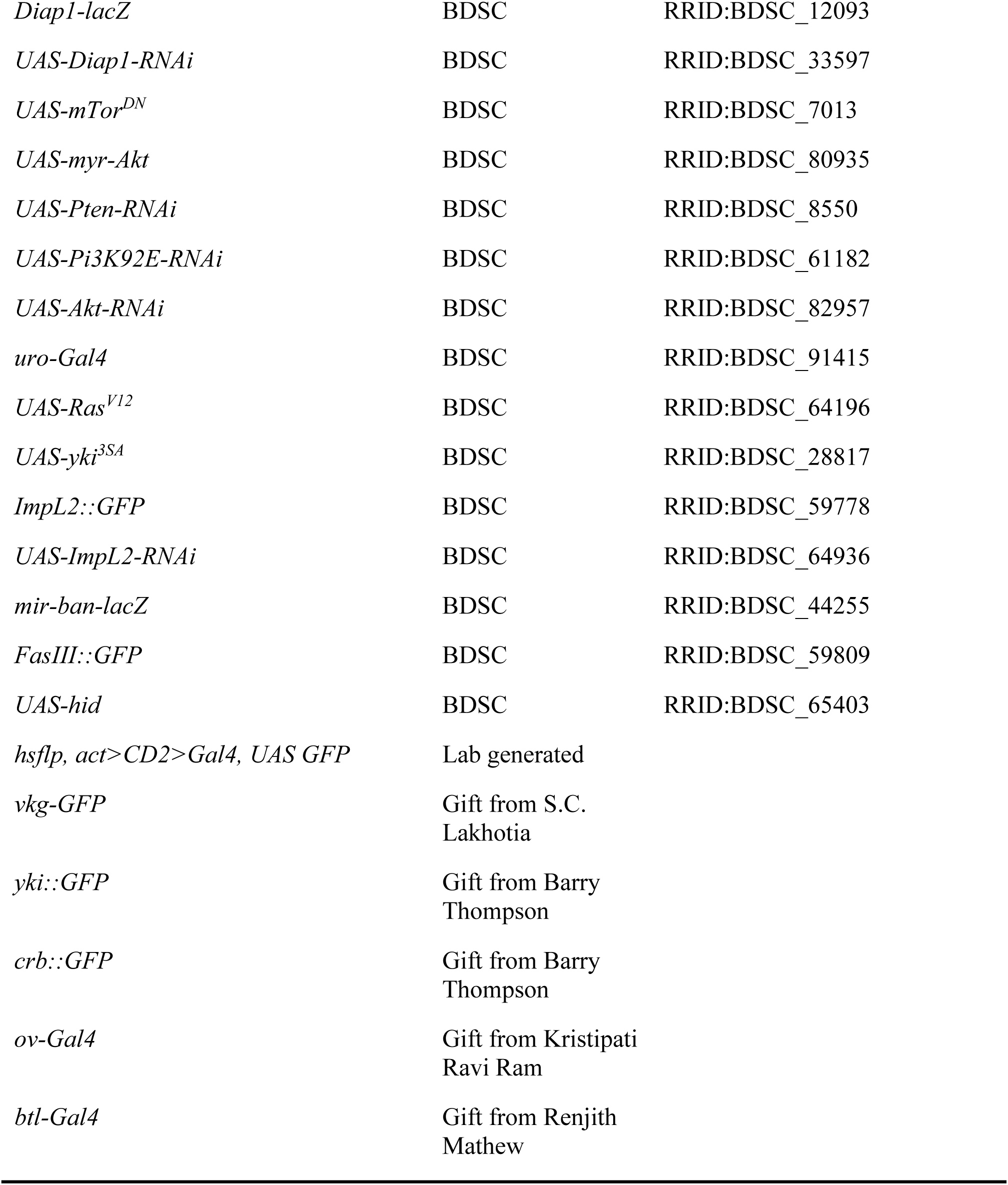

| REAGENTS OR RESOURCE | SOURCE | IDENTIFIER |
| --- | --- | --- |
| **Antibodies** |  |  |
| Mouse anti-FasIII | DSHB | 7G10 |
| Mouse anti-Coracle | DSHB | C566.9 |
| Mouse anti-α-tubulin | DSHB | 12G10 |
| Mouse anti-βPS-Mys | DSHB | CF.6G11 |
| Mouse anti-MMP1, 3, 9 | Santa Cruz<br>Biotechnology | Gift from Ashwani Thakur |
| Mouse β-Gal | Sigma-Aldrich | G6282 |
| Rabbit anti-Lgl | Sigma-Aldrich | Gift from Fumio Matsuzaki |
| Rabbit p-S6K | CST | 9205 |
| Rabbit p-4EBP | CST | 2855 |
| Mouse anti-β-Galactosidase | Sigma | SAB4200805 |
| Alexa Fluor 488 Goat anti-mouse IgG | Invitrogen | A11007 |
| Alexa Fluor 555 Goat anti-mouse IgG | Invitrogen | A32727 |
| Alexa Fluor 633 Goat anti-mouse IgG | Invitrogen | A21050 |
| Alexa Fluor 488 Goat anti-rabbit IgG | Invitrogen | A21206 |
| Alexa Fluor 555 Goat anti-rabbit IgG | Invitrogen | A21428 |
| Alexa Fluor 633 Goat anti-rabbit IgG | Invitrogen | A21070 |
| Nile Red | Sigma | 19123 |
| TO-PRO-3 | Invitrogen | S33025 |
| Phalloidin-555 | Invitrogen | A34055 |
| Vectashield | Vector<br>Laboratories | #H-1000 |
| Software and Algorithms |  |  |
| Leica LAS AF software | Leica microsystem | http://www.leicamicrosystems.com/productes/microscope-software/ |
| Prism | Graph pad | https://www.graphpad.com/ |
| Photoshop | Adobe CS-6 | http://www.adobe.com |
| Illustrator | Adobe CC | http://www.adobe.com |
| BioRender | - | https://biorender.com/ |
| FIJI | ImageJ | https://imagej.net/software/fiji/ |

Other
|  |  |
| --- | --- |
| Leica TCS SP5 microscope | Leica microsystem |
| Leica M205 FA stereomicroscope | Leica microsystem |
| *Drosophila* strains |  |  |
| *UAS-GFP.nls* | BDSC | RRID: BDSC_4776 |
| *tubP-Gal80[ts]* | BDSC | RRID: BDSC_7108 |
| *UAS-yki-RNAi* | BDSC | RRID: BDSC_34067 |
| *UAS-sd-RNAi* | BDSC | RRID: BDSC_55404 |
| *UAS-Hth-RNAi* | BDSC | RRID: BDSC_34637 |
| *UAS-mir-ban-RNAi* | BDSC | RRID:BDSC_60671 |
| *Diap1-lacZ* | BDSC | RRID:BDSC_12093 |
| *UAS-Diap1-RNAi* | BDSC | RRID:BDSC_33597 |
| *UAS-mTor<sup>DN</sup>* | BDSC | RRID:BDSC_7013 |
| *UAS-myr-Akt* | BDSC | RRID:BDSC_80935 |
| *UAS-Pten-RNAi* | BDSC | RRID:BDSC_8550 |
| *UAS-Pi3K92E-RNAi* | BDSC | RRID:BDSC_61182 |
| *UAS-Akt-RNAi* | BDSC | RRID:BDSC_82957 |
| *uro-Gal4* | BDSC | RRID:BDSC_91415 |
| *UAS-Ras<sup>V12</sup>* | BDSC | RRID:BDSC_64196 |
| *UAS-yki<sup>3SA</sup>* | BDSC | RRID:BDSC_28817 |
| *ImpL2::GFP* | BDSC | RRID:BDSC_59778 |
| *UAS-ImpL2-RNAi* | BDSC | RRID:BDSC_64936 |
| *mir-ban-lacZ* | BDSC | RRID:BDSC_44255 |
| *FasIII::GFP* | BDSC | RRID:BDSC_59809 |
| *UAS-hid* | BDSC | RRID:BDSC_65403 |
| *hsflp, act>CD2>Gal4, UAS GFP* | Lab generated |  |
| *vkg-GFP* | Gift from S.C. Lakhotia |  |
| *yki::GFP* | Gift from Barry Thompson |  |
| *crb::GFP* | Gift from Barry Thompson |  |
| *ov-Gal4* | Gift from Kristipati Ravi Ram |  |
| *btl-Gal4* | Gift from Renjith Mathew |  |

## ACKNOWLEDGEMENTS

We would like to thank Barry Thompson, Renjith Mathew, S.C. Lakhotia and Kristipati Ravi Ram for fly stocks, and Ashwani K Thakur for antibodies.

