## Supplementary figures and images for "Hippo effector, Yorkie, is a Tumor Suppressor in Select *Drosophila* Squamous Epithelia"

### Supplementary Figure 1

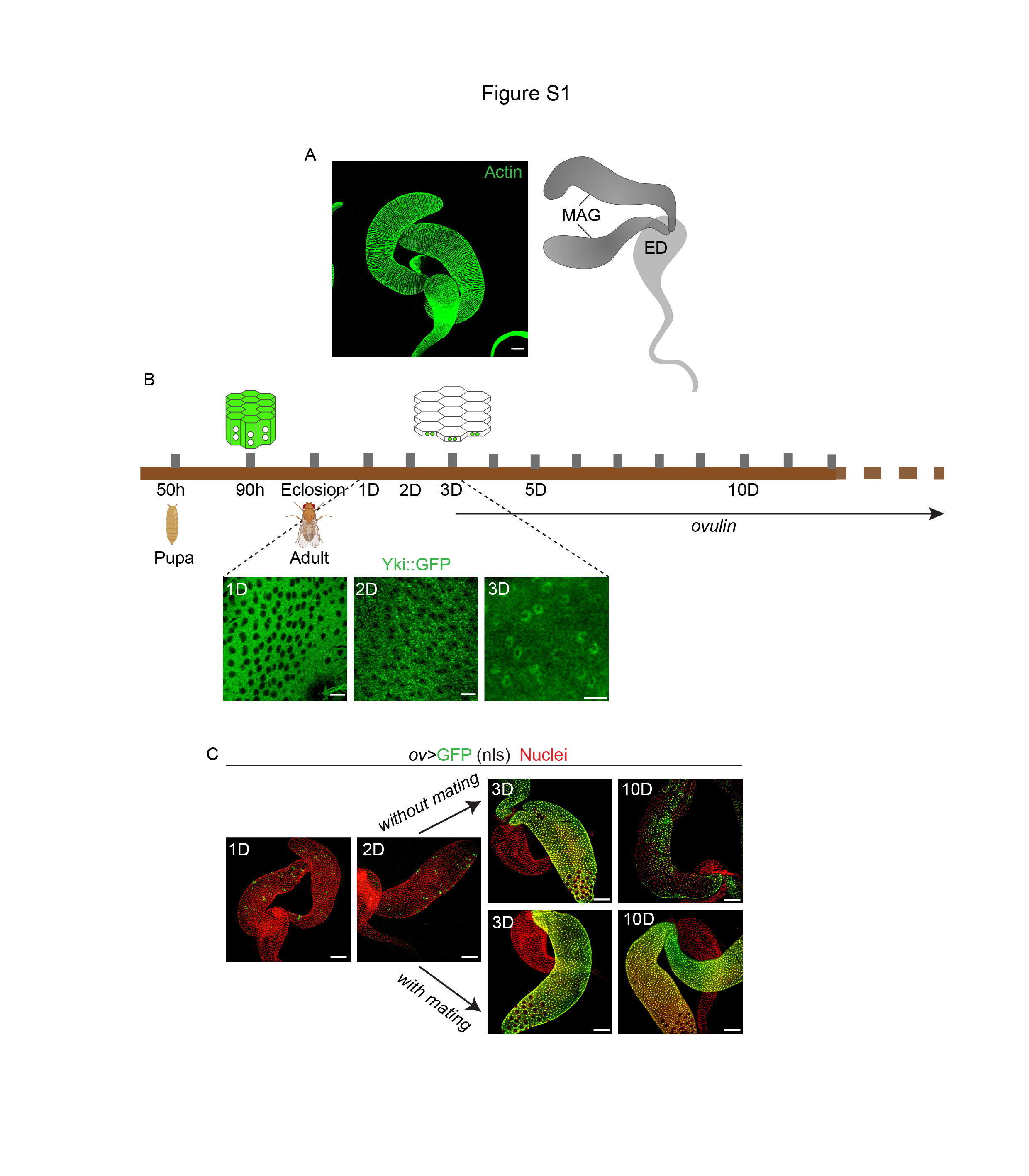

### Supplementary Figure 2

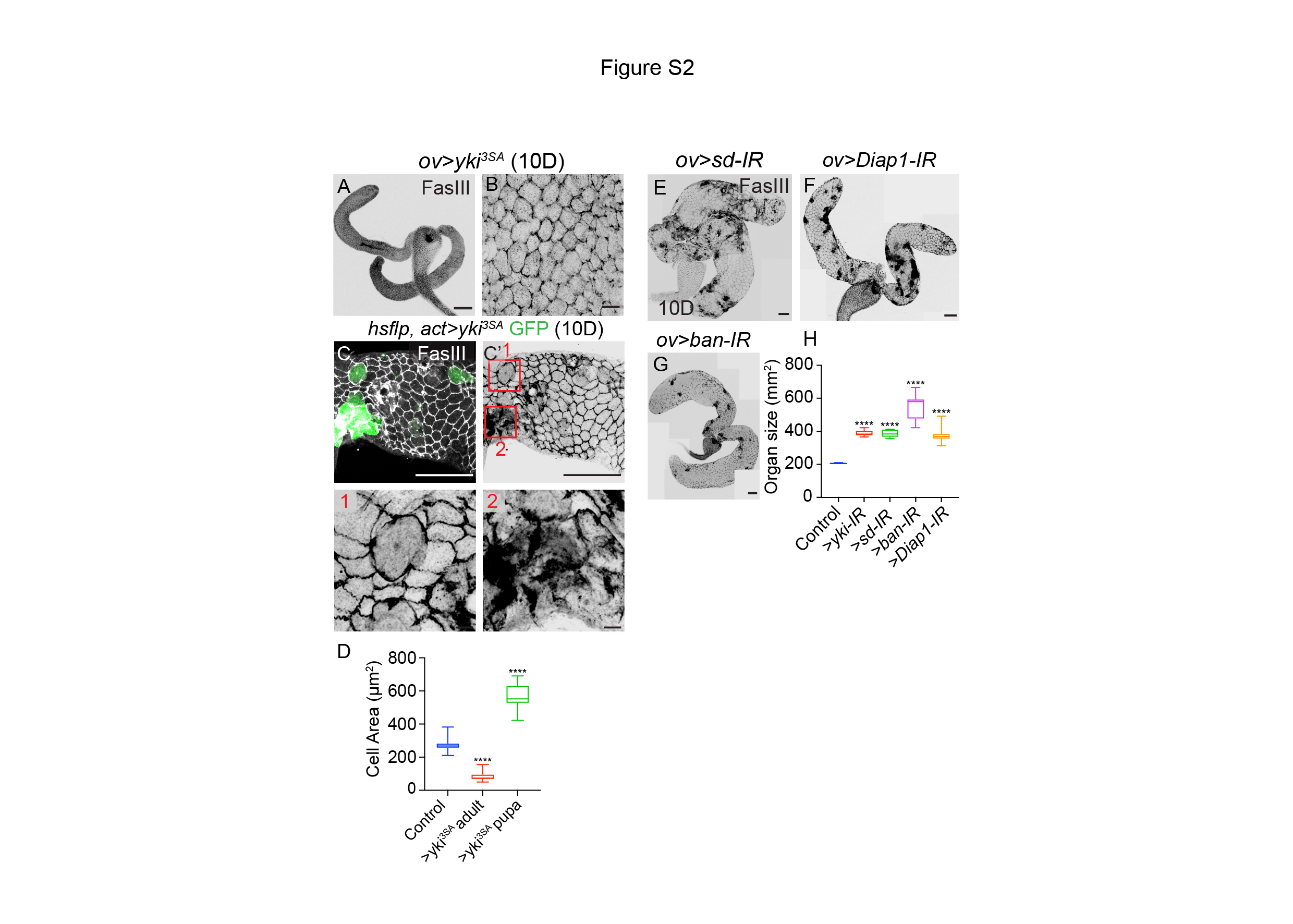

### Supplementary Figure 3

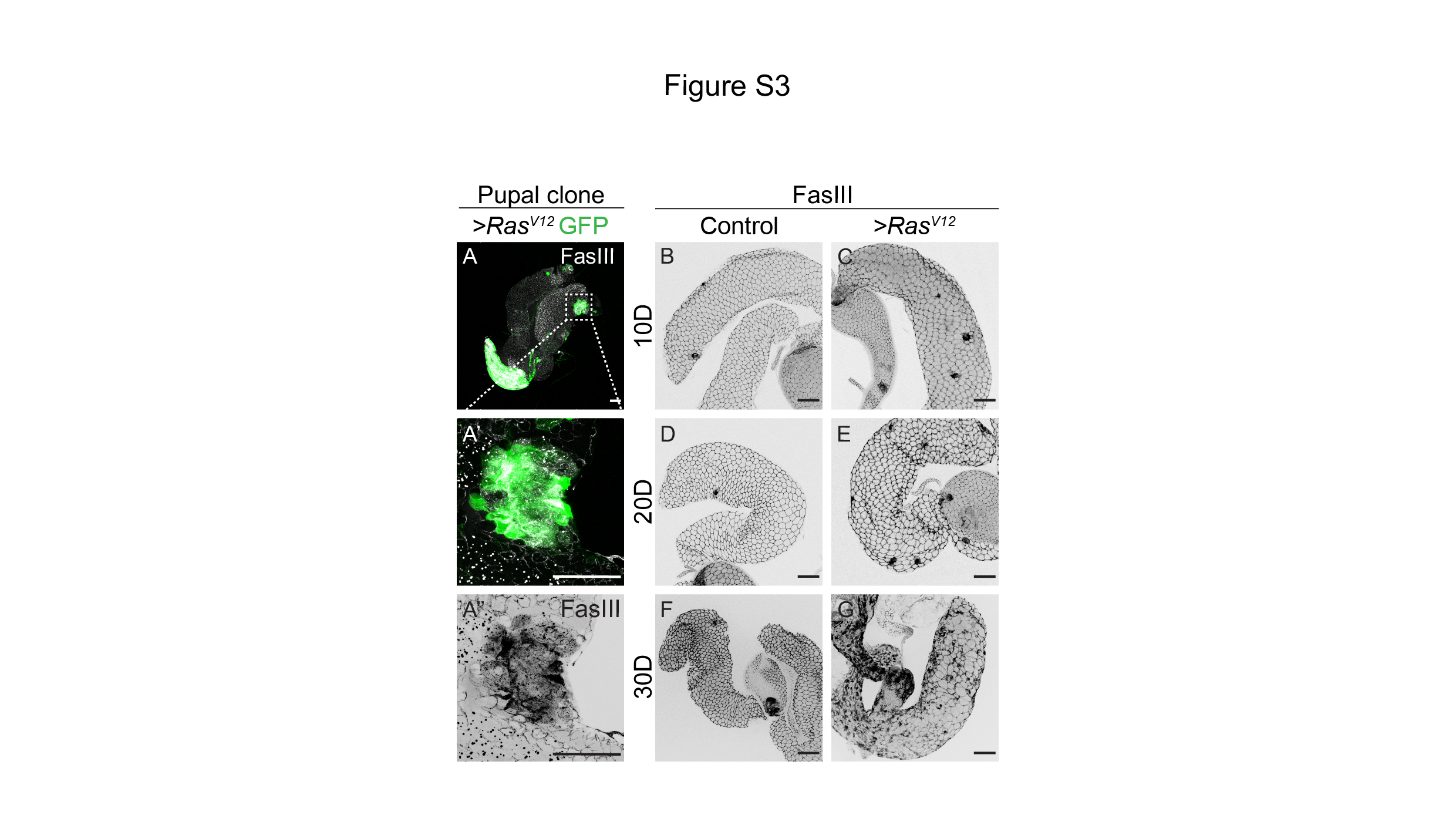

### Supplementary Figure 4

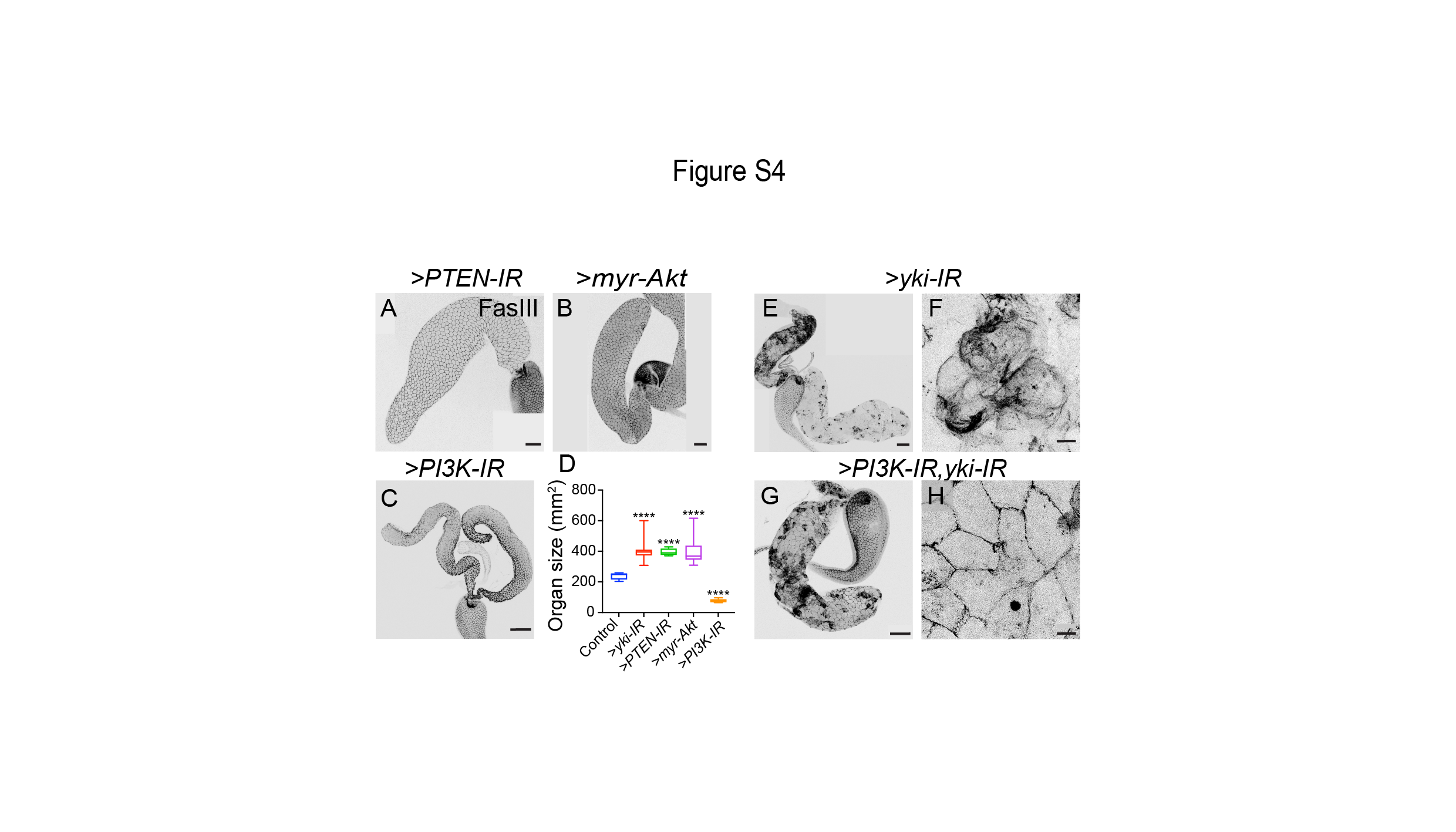

### Supplementary Figure 5

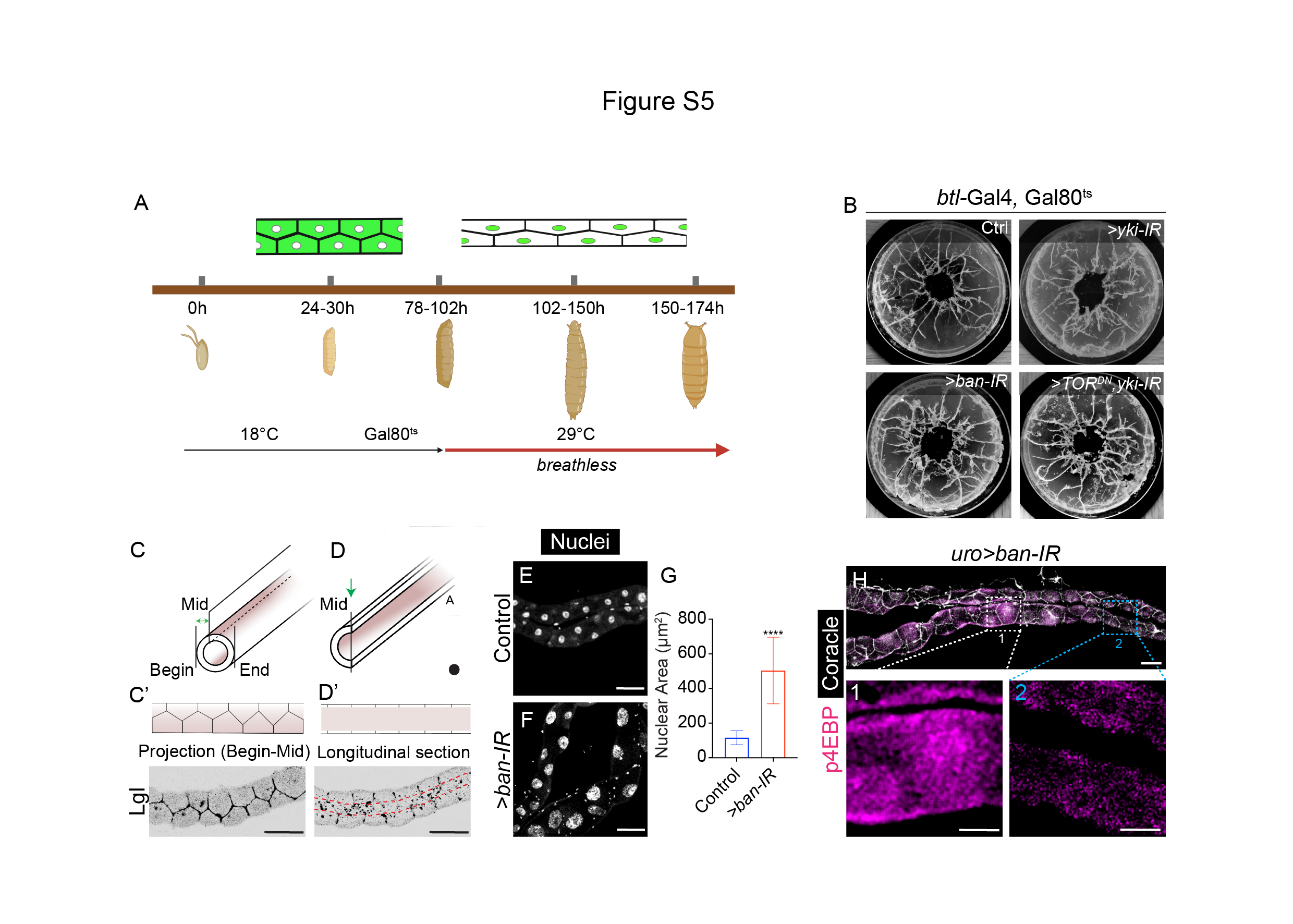
