## Supplementary Figure Legends for "Hippo effector, Yorkie, is a Tumor Suppressor in Select *Drosophila* Squamous Epithelia"

**SUPPLEMENTARY Figure 1-5 legends**

**Fig. S1. Experimental strategy of assessing the role of Yki signaling in the squamous epithelium of adult MAG. (A)** The adult male reproductive system stained for actin (green), displays paired MAG opening into a common ejaculatory duct (ED). The cartoon on the right marks these anatomical features. **(B)** A schematic presentation of Yki::GFP localization in the cytoplasm of one-day-old adult MAG epithelium and its subsequent nuclear translocation in that of a three-day-old adult coinciding with the columnar-to-squamous transition of its epithelium. **(C)** Endogenous *ovulin, ov,* expression, as visualized by the *ov-Gal4>GFP* (green) expression in the adult MAG. *ov* expression is initiated on the third day of adult eclosion, which progressively fades by day ten of adult life when female mates are not added to the culture. By contrast, on the arrival of female mates, robust *ov>GFP* expression is sustained for a far more extended period.

Scale bar: (A) 100 µm, (B) 10 µm (C) 100 µm.

**Fig. S2. Fallouts of gain and loss of Yki signaling in pupal and adult MAG.** (**A-D**) Experimental gain of Yki signaling in adult squamous epithelium MAG (*ov>*$yki^{3SA}$) reduces its organ size (A, B). By contrast, a gain of Yki signaling in somatic clones induced in pupal MAG (50h) (*hs-flp, act>*$yki^{3SA}$, GFP, C, C') induces cell hypertrophy (box 1) and neoplastic transformations (box 2). Magnified views of Box 1 and Box 2 are presented in the bottom panel. (D) Quantifying adult MAG cell area upon the gain of Yki signaling in pupa and adult. While the pupal gain of Yki signaling induces cell hypertrophy, the converse is the fallout upon its gain in adult squamous epithelium. Unpaired *t*-test of average cell area between control and Yki gain (****$p$<0.0001). Data are represented as mean ± *SD* ($n$ = 50 for control (*ov>+*) and *ov>*$yki^{3SA}$, $n$ =10 for *hs-flp, act>* $yki^{3SA}$).

(**E-H**) Whole mounts of adult MAG displaying knockdown of *yki* partners, the transcription factor *sd* (E), and their downstream targets *Diap1* (F) and *bantam* (G) under the *ov-Gal4* driver. (H) Quantification of MAG size of the indicated genotypes. Unpaired *t*-test of average cell area between control and those displaying knockdown of different components of Yki signaling pathway reveal ****$p$<0.0001). Data represent mean ± *SD* ($n$=10 for each indicated genotype).

Scale bar: (A) 100 µm, (B) 10 µm, (C, C') 100 µm, (box 1 and 2) 10 µm, (E-G) 100 µm.

**Fig. S3**. **$Ras^{V12}$ gain is oncogenic in the pupal MAG's columnar epithelium but not in its adult squamous counterpart. (A)** $Ras^{V12}$-expressing somatic clones (GFP, green) induced in the pupa (50h) and examined in 5-day-old adult MAG after FasIII immunostaining (grey/white). The bottom two panels (A', A") display a higher magnification of the boxed area. Note the tumor cell morphology, corresponding to the GFP-marked $Ras^{V12}$ clone, marked by loss of septate junction-specific organized FasIII staining (A"). (**B-G**) Low magnification view of entire MAG immunostained for FasIII (grey) of control and $Ras^{V12}$-overexpressing MAG; age of the host adults is indicated. With progressive aging, $Ras^{V12}$-expressing adult MAG displays altered cytoarchitecture and delamination (see Fig. 4) but does not turn into SCC.

Scale bars: (A-G) 100 µm.

**Fig. S4. Inhibition of TOR signaling rescues *yki* loss-driven MAG-SCC.** (**A-D**) Comparison of MAG organ sizes of *ov>PTEN-IR* (A), *ov>myr-Akt* (B), *ov>PI3K-IR* (C) and its quantification (D). Unpaired *t*-test of average cell area between control and test genotypes display ****$p$<0.0001. Data display mean ± *SD* ($n$=10 for each indicated genotype). **(E-H)** By contrast to *yki* knockdown-induced MAG-SCC (E-F), a simultaneous knockdown of PI3K signaling (*ov>PI3K-IR, yki-IR,* G-H) arrested MAG SCC marked by restoration of FasIII-marked cell outlines (grey).

Scale bars: (A-E, G) 100 µm, (F, H) 10 µm.

**Fig. S5. *yki* loss upregulates TOR signaling in the squamous epithelium of tracheal trunks and Malpighian tubules. (A)** Schematic display of Yki::GFP expression in the dorsal trunks of the trachea network reveals its cytoplasmic localization till the second instar, which becomes nuclear during the third larval instar. Expression of the *btl* is initiated during embryonic tracheal development and lasts throughout the larval development (Samakovlis et al., 1996). To achieve a selective, third larval instar-specific expression of the *btl-Gal4* driver in the dorsal tracheal trunks—post-nuclear Yki translocation—we coexpressed a temperature-sensitive Gal-80$^{ts}$ driver (McGuire et al., 2003) in larval cultures grown at 18$^{o}$C. A shift in larval growth temperature to 29$^{o}$C starting the third larval instar—and a consequent inactivation of the Gal-80$^{ts}$ driver led to a *btl-Gal4*-driven expression of the *yki-IR* or other transgenes. **(B)** Control and Yki signaling comprised larvae displayed comparable burrowing-tunneling, revealing that knockdown of Yki signaling at this stage does not induce hypoxia (Qiang et al., 2018).

(**C**) Schematic representation of a Malphigian tubule, wherein longitudinal optical sections are projected from the beginning to the mid position (green double arrowhead). In such a scenario, (**C'**) flattened Malphigian tubule cells wrapped around the tubule are seen to occupy half the circumference of the tube, sharing their cell margins in the center. The bottom panel shows actual projections Lgl marked (marking lateral epithelial membrane, grey), revealing this scenario mentioned in (C, C') above. **(D)** By contrast, in an optical section passing through the midline of a radially symmetric tube (**D'**), the margin of the lateral membranes of the cells wrapped around the tubule are seen to flank the central lumen. The bottom panel shows an actual single optical section along the midline of the Malphigian tubule immunostained for Lgl (grey). Broken lines highlight the lumen.

**(E-G)** Malpighian tubules of control (E) and *uro>ban-IR* (F), stained with TO-PRO, white. Note the enlargement of the nuclear size in the latter. (G) Quantification of the nuclear size of indicated genotype. Unpaired $t$-test of average nuclear size between control and *ban* loss displays ****$p$<0.0001. Data are presented as mean ± *SD* ($n$=50 for each genotype).

**(H)** Gain of the TOR signaling target, 4EBP-P (magenta, box 1), in the *uro>ban-IR*-expressing region in the main segment of adult Malpighian tubule, unlike that of the non-*uro>ban-IR*-expressing parts of the tubule (box 2). A septate junction marker, Coracle (white), marks the cell boundaries (Lamb et al., 1998).

Scale bars: (C-F) 50 µm, (H) 100 µm, (Box 1 and 2) 50 µm.
